# Transcriptional natural variation at *FLM* induces synergistic pleiotropy in *Arabidopsis thaliana*

**DOI:** 10.1101/658013

**Authors:** Mathieu Hanemian, François Vasseur, Elodie Marchadier, Elodie Gilbault, Justine Bresson, Isabelle Gy, Cyrille Violle, Olivier Loudet

**Affiliations:** Institut Jean-Pierre Bourgin, INRA, AgroParisTech, CNRS, Université Paris-Saclay, 78000, Versailles, France; CEFE, CNRS, Univ Montpellier, Univ Paul Valéry Montpellier 3, EPHE, IRD, F-34090 Montpellier, France; Laboratoire d’Ecophysiologie des Plantes sous Stress Environnementaux (LEPSE), INRA, Montpellier SupAgro, UMR759, F-34060 Montpellier, France

**Keywords:** Alternative splicing, Natural variation, Pleiotropy, Leaf economics spectrum, Flowering time

## Abstract

Investigating the evolution of complex phenotypes and the underlying molecular bases of their variation is critical to understand how organisms adapt to the environment. We used leaf growth as a model trait as it is highly integrative of internal and external cues and relies on functions at different levels of the plant organization. Applying classical quantitative genetics on a recombinant inbred line population derived from a Can-0 x Col-0 cross, we identified the MADS-box transcription factor *FLOWERING LOCUS M* (*FLM*) as a player of the phenotypic variation for leaf growth and colour. Interestingly, we showed that allelic variation at *FLM* modulates plant growth strategy along the leaf economics spectrum, a trade-off between resource acquisition and resource conservation observable across thousands of plant species. We demonstrated that the functional differences at *FLM* relies on a single intronic substitution, disturbing transcript splicing and leading to a low expression of the active *FLM* transcript. Using phenotypic and climatic data across Arabidopsis natural populations, our work shows how noncoding genetic variation of a single gene may be adaptive through synergistic pleiotropy.

## INTRODUCTION

Evolution is a continuous and complex process requiring the coordinated changes in many traits according to environmental selective pressure, resulting in an optimal fitness in a given habitat at a given time point. Owing to the numerous traits that need to evolve coordinately, mathematical models predicted that complex organisms would evolve more slowly than simple organisms towards a fitness optimum when considering mutations of same effect size (Fisher, 1930; Orr, 2006). This was called the “cost of complexity” or the “cost of pleiotropy” since those models assumed that every gene is able to affect every trait (universal pleiotropy).

Pleiotropy can have several meanings but overall states that one variant or gene controls several phenotypes (Paaby and Rockman, 2013). The nature of pleiotropy strongly depends on the traits measured (molecular, physiological, metabolic…) and the level of organization considered (from cell to population). Moreover, one need to be cautious about the relevance of the observed pleiotropy according to the method used to assess it. For instance, studying knock-out mutants may not reflect the diversity of mutational effects selected in nature. Also, when classical quantitative genetics is used to study pleiotropy, the main question is to distinguish real pleiotropy from genetic linkage, i.e. several independent quantitative trait loci (QTLs) in the same region controlling different traits, because mapping QTL in segregating populations has limited resolution (Marchadier et al., 2019). Finally, many related traits may be correlated but may just represent a cause-and-effect relationship, in other word an indirect effect.

Using reverse genetics and QTL mapping approaches in various organisms, it has been observed that pleiotropy is relatively rare (Wagner et al., 2008; Wang et al., 2010). Moreover, a high degree of pleiotropy (i.e. the diversity of traits controlled by one variant/gene) is thought to limit the potential of adaptation because it would be more likely to affect negatively some functions while improving others (‘antagonistic pleiotropy’), thus changing the optimal balance selected over the course of evolution. From this perspective, adaptation speed would rather depend on the degree and the modularity of pleiotropy (the combination of traits controlled by one variant/gene) than on the complexity of organisms (Wagner and Zhang, 2011; Wagner et al., 2007). Inspection of phenotypic data from yeast, nematode and mice mutants revealed that an intermediate degree of pleiotropy seems the best trade-off to reach the highest rate of adaptation (Wang et al., 2010). A recent study revealed that indeed, a combination of QTLs having an intermediate degree of pleiotropy, with QTLs affecting specifically one trait were able to promote a fast adaptation (Frachon et al., 2017).

In plants, phenotypic traits involved in the strategies for resource acquisition and use, called functional traits, have been widely investigated across species (Violle et al., 2007; Díaz et al., 2016). Several trait-environment relationships have been identified and are assumed to be representative of species adaptation to various environmental conditions. Ecologists have long recognized that functional traits are often correlated with each other, and that adaptation to new environments requires a simultaneous change in these traits due to resources limitation and biophysical constraints (Reich, 2014). For instance, the rate of carbon fixation through photosynthesis is linked to the structure of the leaves, such as specific leaf area (SLA, i.e. the ratio of leaf area over leaf dry mass) and leaf dry matter content. Leaf structure and growth is also related to leaf chemical composition and its lifespan (Wright et al., 2004; Tisné et al., 2010). Together, these traits shape a trade-off between resource acquisition and resource conservation, observable across thousands of plant species and coined under the term “Leaf Economics Spectrum” (LES) (Wright et al., 2004). At one end of the spectrum are plants with resource-conservative strategies, characterized by long-lived, tough leaves with low nutrient concentration and low net photosynthetic rate. At the other end are plants with resource-acquisitive strategies, characterized by short-lived, flimsy leaves with high nutrient concentration and high net photosynthetic rate. The same pattern of phenotypic variability has been observed within the model species *Arabidopsis thaliana*, where the LES is related to life history traits’ variation between accessions such as flowering time (Vasseur et al., 2012, 2018).

While the correlation between traits and their functions in various environment starts to be well-characterized (Garnier et al., 2017), we do not know so much about their genetic architecture and molecular bases, as well as the importance of pleiotropy in the evolution of organisms. Inference about natural selection and adaptive processes at the origin of phenotypic diversification along environmental gradients indeed requires the examination of the molecular determinants of trait variation within species and across populations (Donovan et al., 2011).

Synergistic evolution of multiple traits can be achieved by multiple genes (polygenic hypothesis) with medium to small effect, and/or by few genes with multiple phenotypic effects (pleiotropy hypothesis) (Pritchard et al., 2010; Wagner and Zhang, 2011). While the former might be slow but fine-tunable, the latter is expected to play a predominant role for adaptation to abrupt environmental changes with a bet-hedging and risky evolutionary path, as a single locus would strongly modify plant ecological strategy (Orr, 2005). Dissecting the genetic architecture and identifying the genes behind correlated traits would help to assess the number of loci involved, their individual effects, their degree of pleiotropy as well as their potential for adaptation.

Examples of genes responsible for the natural variation of single traits such as salinity tolerance (Baxter et al., 2010), flowering time (Stinchcombe et al., 2004; Liu et al., 2014), seed dormancy (Kronholm et al., 2012; Vidigal et al., 2016) or herbivore resistance (Brachi et al., 2015) are increasing. However, these studies are about a specific type of adaptation essentially having a simple genetic architecture at the species scale, i.e. with one or a few genes controlling most of the phenotypic variation. Studies aiming at the dissection of the genetic bases of the phenotypic integration of several complex traits such as plant growth or fitness are more scarce and rarely reach the stage of the functional validation of the candidate genes (Vasseur et al., 2012, 2014; Ågren and Schemske, 2012; Postma and Ågren, 2016).

Major developmental switches during plant growth cycle interact with plant physiology and fitness, among which the timing to flowering is expected to play a crucial adaptive role to ensure plant reproductive success. Some flowering time genes were shown to have pleiotropic functions during development. Consistently, a previous study has shown tight relationships between flowering time and the vegetative growth dynamics in *A. thaliana* (Bac-Molenaar et al., 2016). In *A. thaliana*, the molecular control of flowering time is orchestrated by a complex network of genes integrating several signals that converge towards a few regulators (Blümel et al., 2015; Cho et al., 2017). Among them, FLOWERING LOCUS C (FLC) is a MADS-box transcription factor known as a repressor of flowering (Michaels and Amasino, 1999; Lee et al., 2000). FLC expression is down-regulated by extended cold exposure to prime winter-annual *A. thaliana* populations for flowering in spring (Whittaker and Dean, 2017). Interestingly, FLC also plays a role in germination time, another critical phase in plant development (Chiang et al., 2009; Chen and Penfield, 2018) as well as vegetative development (Willmann and Poethig, 2011). Furthermore, FLC was shown to bind to promoters of hundreds of genes involved in stress-related and hormonal pathways, activating or repressing their expression (Deng et al., 2011).

Likewise, the role of pleiotropy in drought adaptation was studied by assessing physiological traits that are correlated in nature such as water use efficiency (WUE) and flowering time (McKay et al., 2003). It was later shown that natural variation of *FRIGIDA*, another major regulator of flowering time which activates *FLC* expression, modulates drought tolerance strategy through a pleiotropy exerted on several physiological traits (Lovell et al., 2013). Another example is that of *ERECTA* and its effect on functions like transpiration efficiency as well as inflorescence development (Masle et al., 2005). To understand how phenotypic integration regulates plant adaptation to contrasted environments, it is crucial to examine the genetic underpinnings of relationships between life history, leaf physiology and climate.

Here, we sought to decompose complex traits into individual genetic and molecular components and have thus identified the molecular basis of a QTL controlling both vegetative growth and time to flowering in *A. thaliana.* Through a combination of quantitative genetics, molecular biology, ecophysiology and population genetics, we show how modulating the transcriptional output of a gene can have a pleiotropic effect on multiple functions affecting life history and plant physiology. These results are exploited to distinguish pleiotropy from linkage in a concrete case, and to connect general theories about adaptation and pleiotropy.

## RESULTS

### From several QTLs to a pleiotropic gene

Using the *Phenoscope* phenotyping platform for *A. thaliana*, we measured projected rosette area 29 days after sowing (PRA29) during the vegetative phase of growth on 360 individuals of a Recombinant Inbred Line (RIL) population, derived from the cross between Can-0 and Col-0. We performed a QTL analysis to assess the genetic bases of variation in plant growth. In addition, several other image-derived traits such as leaf colour, a surrogate for pigment content, were investigated (Sass et al., 2012). QTLs for several of the measured traits co-localize at the bottom of chromosome 1 with mild effect on growth and large effect on leaf colour (Fig. S1A). We then followed a classical fine-mapping approach to identify the gene(s) underlying these trait variations. Thus, we selected the F7 RIL line 19RV337 with a residual heterozygous region covering the expected locus to build the Heterogeneous Inbred Family (HIF) 19HV337 by selecting progenies homozygous for each of the parental allele at the candidate region. The phenotypic comparison of homozygous siblings confirmed that the allele originating from Can-0 (*Can*) was promoting PRA29 with respect to the allele originating from Col-0 (*Col*), and also influenced leaf colour (Fig. S1B). Phenotyping 12 selected recombinant HIF lines (rHIF) derived from 19HV337 confirmed the association of phenotypic variation with the candidate region containing 10 genes (Fig. S1C). The Can allele at this locus conferring both higher vegetative growth and earlier bolting with respect to the Col allele, we focused on the candidate gene coding for *FLOWERING LOCUS M* (*FLM*). Compared to its wild type (WT) control, the mutant *flm-3* displays similar phenotypic effects on leaf growth and colour as rHIF099-Can compared to rHIF099-Col (our ultimate rHIF segregating at *FLM*), indicating that Can-0 could potentially harbour a hypofunctional *FLM* allele (Fig. 1A).

**Figure 1:**
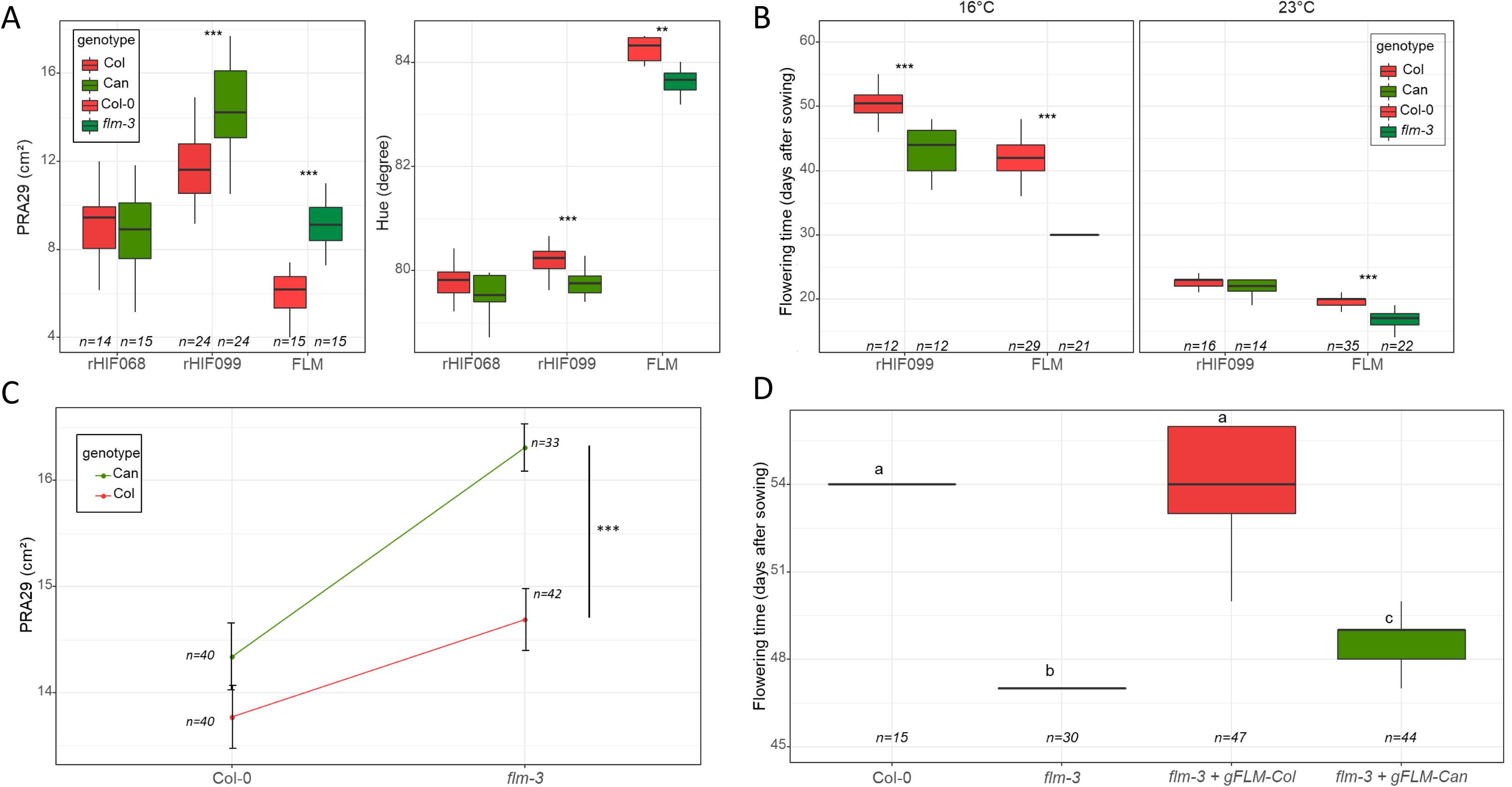
Natural variation at *FLM* is responsible for the QTLs observed at the end of chromosome 1. A. Boxplots comparing PRA29 and Hue, in the two most informative recombinant lines (rHIF068 is fixed at *FLM* and rHIF099 is segregating at *FLM*), and in *flm-3* versus wild-type allele (Col-0 background). B. Modulation of flowering time according to the growth conditions at 16°C and 23°C (in long days) in both rHIF099 lines and in *flm-3* versus wild-type allele (Col-0 background). C. Quantitative complementation assay assessing the effect on PRA29 of each of the possible genotypic combination between the Can or Col alleles at the QTL (in the rHIF background) and the wild-type or mutant (*flm-3*) alleles at FLM (in the Col-0 background). PRA29 was obtained using the *Phenoscope*. Error bars indicate the standard deviation of the mean. D. Functional complementation based on flowering time of the *flm-3* mutant transformed with the genomic fragments of *FLM* (*gFLM*) from Col-0 or Can-0.

FLM is a MADS-box transcription factor inhibiting flowering at cool temperature and whose repressive action is alleviated when temperature increases moderately (Scortecci et al., 2001; Balasubramanian et al., 2006). We showed that flowering time was indeed segregating specifically at cool temperature in rHIF099, similarly to the difference observed between *flm-3* and its WT counterpart (Fig. 1B). To further confirm *FLM* as the gene underlying the vegetative growth-related QTL, we performed a quantitative complementation assay (Mackay, 2004) relying on the comparison, in F1 hybrid plants, of the *Can* and *Col* alleles with respect to the *flm-3* allele. Hybrids were derived from crosses between rHIF099 (segregating at *FLM*) and *flm-3* or its WT background (Fig. 1C). Highly significant differences in plant growth were observed when *flm-3* was complemented by either of the QTL alleles (p=2.638e-05). Although the significance of the interaction between the QTL allele and the mutant genotype was itself only suggestive (p=0.069), this overall indicates that the *Can* allele at *FLM* is less able to complement *flm-3* than the *Col* allele (Fig. 1C). Finally, we genetically transformed the *flm-3* mutant with a genomic fragment containing 3kb of the promoter and the full *FLM* genomic region (including introns and the 3’UTR) from either *FLM-Can* or *FLM-Col. FLM-Col* fully complemented *flm-3* to WT level in terms of flowering time, unlike *FLM-Can*, confirming that this gene is hypofunctional in Can-0 (Fig. 1D). Altogether, these data show that allelic variation at *FLM* is underlying the leaf growth and colour QTLs segregating in Can-0 x Col-0 at the lower end of chromosome 1.

### Molecular bases of the variation at *FLM*

De novo sequencing of PCR-amplified *FLM-Can* and *FLM-Col* alleles did not show any polymorphism within the coding sequence between these alleles (SupFile1 and Fig. S2). We therefore assessed expression of the *FLM* alleles using 4 weeks-old rosettes of the rHIF099 segregating for *FLM. FLM*-dependent thermosensitivity of flowering has been proposed to rely on a differential alternative splicing promoting either the production of the repressive isoform *FLM*-*β* at cool temperature or *FLM-δ* when temperature increases (Posé et al., 2013). Semi-quantitative PCR showed that *FLM-β* transcript accumulation is strongly reduced in rHIF099-Can compared to rHIF099-Col (Fig. S3A & S3B). The result on *FLM-δ* was less clear due to amplification of several other isoforms (see below). Using qRT-PCR, we quantified *FLM-β* expression as well as the expression level of the 1st exon, which is common to both isoforms and used it as a readout for *FLM* overall transcription (Fig. 2A & 2B). Interestingly, while the expression level of the 1st exon was reduced ∼3 fold comparing Can and Col alleles, *FLM-β* expression was reduced by a factor of ∼29. These data suggest that a mechanism is more specifically reducing the level of the *FLM-β-Can* transcript, although the promoter may also slightly contribute to its decrease as suggested by the relatively moderate reduction of the global expression of the 1st exon. Since *FLM-β* is the active isoform shown to repress flowering time in natural strains of *A. thaliana* (Lutz et al., 2015, 2017; Capovilla et al., 2017), we hypothesized that its differential expression between the Col and Can alleles is causing the phenotypic differences observed.

**Figure 2:**
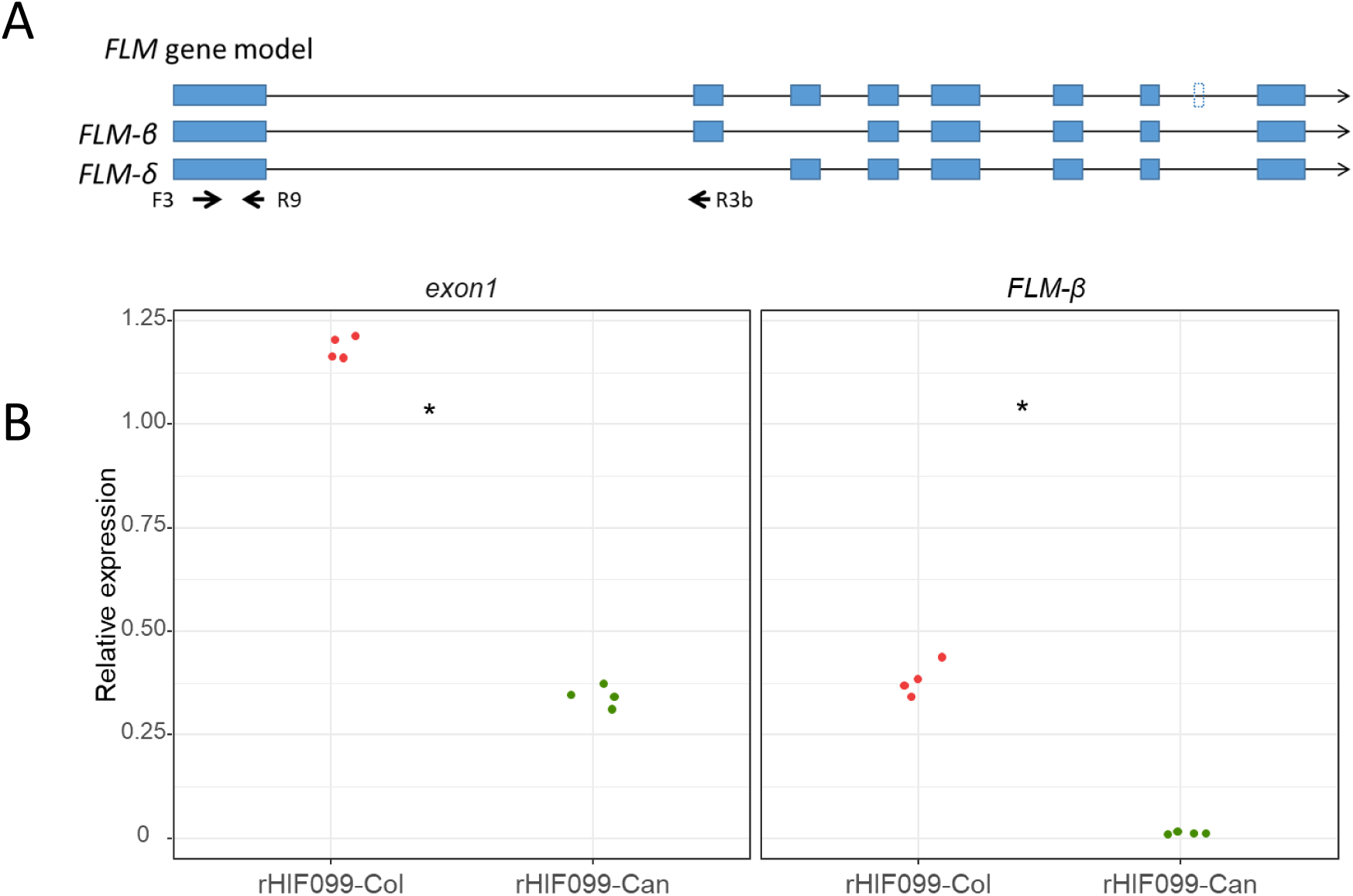
Analysis of *FLM* expression. A. Graphic representation of the *FLM* gene model and transcript as well as its two main splicing variants *FLM-β* and *FLM-δ*. The primers used for quantitative RT-PCR are depicted. B. Transcript accumulation level according to the exon 1 (primers F3 + R9) and *FLM-β* (primers F3 + R3b) were measured by qRT-PCR in rosette leaves grown on the *Phenoscope* from rHIF099 homozygous for the Can or the Col allele. Each dot represents the mean of a technical duplicate of an individual plant.

Another experiment was performed on 4 weeks-old rosettes of rHIF099-Can, rHIF099-Col, Col-0, Can-0 and *flm-3* grown at either 16°C or 23°C. Again, we observed that the expression level of *FLM-β* was strongly reduced in Can-0 compared to Col-0 (∼11 fold and ∼68 fold respectively at 16°C and 23°C), relative to the expression level of the 1st exon (∼3 fold; Fig. S4). These data showed that *FLM-β-Can* is responsive to temperature change but reach a level close to zero at 23°C. Comparable expression levels were observed in wild strains and in the recombinant line for each allele, showing that the genetic background does not affect the differential allelic expression of *FLM*. Thus, we decided to study *FLM* expression using natural strains in the next experiments.

Since we could not amplify specifically the *FLM-δ* isoform, we designed other couple(s) of primers spanning introns and amplifying from the 3rd exon towards the beginning or the end of the gene to perform semi-quantitative PCR (Fig. S3A). The 1st new couple of primers brought similar result as previously, i.e. *FLM-δ* band of expected size seems slightly more intense in Col-0 than in Can-0 but an opposite pattern was observed for the non-specific band of higher molecular weight (Fig. S3C). Another couple of primers was also nonspecific to *FLM-δ* as 2 bands were amplified in both accessions. However, we noticed a size shift of the smaller band (expected size in Col-0) in Can-0 toward a slightly larger molecular weight (Fig. S3C). We thus set out to clone the isoforms present in our samples grown at 23°C to know whether they encode functional proteins and to investigate the *FLM-δ* size shift observed in Can-0. In Col-0, we found a majority of *FLM-β* isoform (34 out of 52 clones sequenced) as previously reported (Sureshkumar et al., 2016) and discovered 2 new isoforms including one which is potentially functional (Fig. S5). In Can-0, we observed that most of the isoforms produced (26/27) contain an insertion of 17 bp before the 3rd exon which explains the size shift observed in the semi-quantitative PCR (Fig. S5). Interestingly, a polymorphism changing a “GG” to an “AG” is located right before this additional stretch of DNA in Can-0 at the position 28,958,437. We deduced that this substitution in Can-0 (hereafter named ‘SNP28958’) is creating a consensus sequence of a splice acceptor site, which modifies *FLM* splicing leading to the incorporation of these additional 17 nucleotides and producing a premature stop codon in the 3rd exon. We showed that SNP28958 abolishes *FLM* function in term of flowering time, PRA29 and leaf colour as the *flm-3* mutant was not complemented by an *FLM-Col* construct substituted with this SNP (Fig. S6A). We detected the *FLM-δ* size shift in these transgenics demonstrating that this single substitution is indeed creating a hot acceptor splice site which generates non-functional *FLM* isoforms (Fig. S6B).

### Pleiotropic effects of *FLM* on leaf physiology

We then wanted to describe how *FLM-Can* and *flm-3* alleles were affecting growth in the longer term under long day photoperiod, by comparison to the Col allele, using parallel PRA and leaf number measurements. As expected, we observed that both weak alleles promote PRA in a first phase of vegetative growth. However, the growth of all lines is slowed down and then stopped maximum one week after bolting, which means that the lines carrying a functional allele of *FLM* continue to grow over a longer period of time to reach the same (if not higher) final rosette sizes (Fig. 3 and Fig. S7). Leaf number was rather similar in a first phase but lines carrying a functional *FLM* allele progressively developed more rosette leaves (Fig. 3 and Fig. S7).

**Figure 3:**
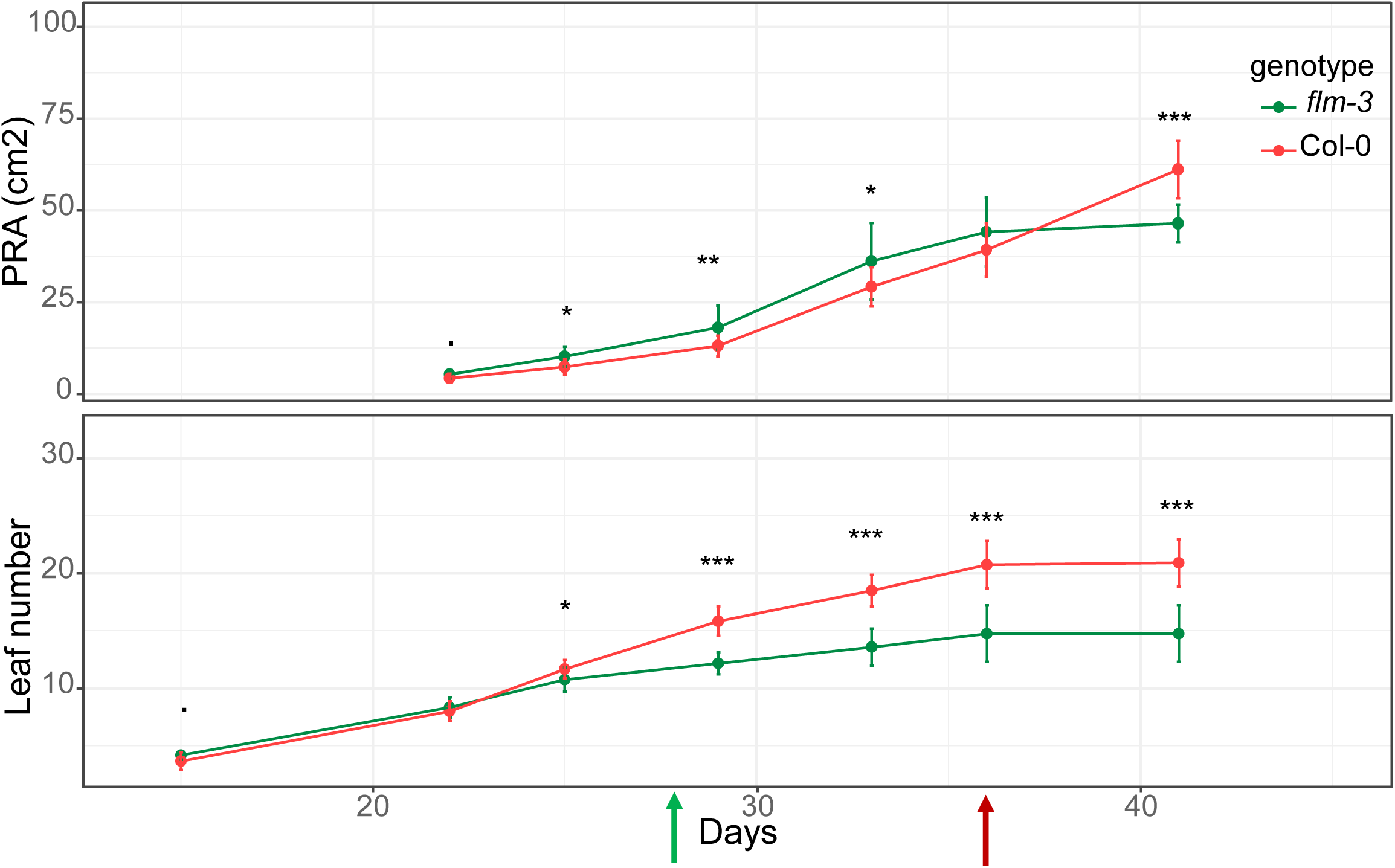
Relationships between vegetative growth and flowering time according to the *FLM* allele. Dynamics of projected rosette area (PRA) and leaf number followed until 41 days after sowing. Green and red arrow indicate the bolting time (floral stem 1cm) in *flm-3* and Col-0 respectively.

As *FLM* affect leaf growth and leaf colour, we hypothesized that it might modulate leaf physiology through pleiotropic effects during vegetative growth. To test this hypothesis, we measured several traits related to leaf physiology and resource-use across a set of segregating lines and *flm* mutants. Leaf temperature measurements showed that leaves of rHIF099-Can and *flm-3* alleles are cooler compared to their respective controls, indicating a higher water flux through the plant and thus a lower WUE (Fig. S8). We also measured net photosynthetic rate per unit dry mass (*A*_mass_) and leaf dry mass per area (LMA), two core traits underlying variations in leaf physiology across and within species. We observed that both *flm-3* and *FLM-Can* triggered a negative correlation between these two traits, similar to the relationships observed across species and within *A. thaliana* natural accessions (Fig. 4) (Kattge et al., 2011; Sartori et al., 2018). The leaf physiology of *flm-3* and *FLM-Can* alleles is indeed shifted toward more acquisitive strategies, *i.e*. a higher *A*_mass_ and a lower LMA. Moreover, relationships slopes were not significantly different (*p* > 0.05) between *flm-3* mutants / WT (SMA slope = -1.80, 95% CI = [-2.11;-1.54], *r*^2^ = 0.17) and among *A. thalian*a accessions (SMA slope = -2.04, 95% CI = [-2.16;-1.92], *r*^2^ = 0.54).

**Figure 4:**
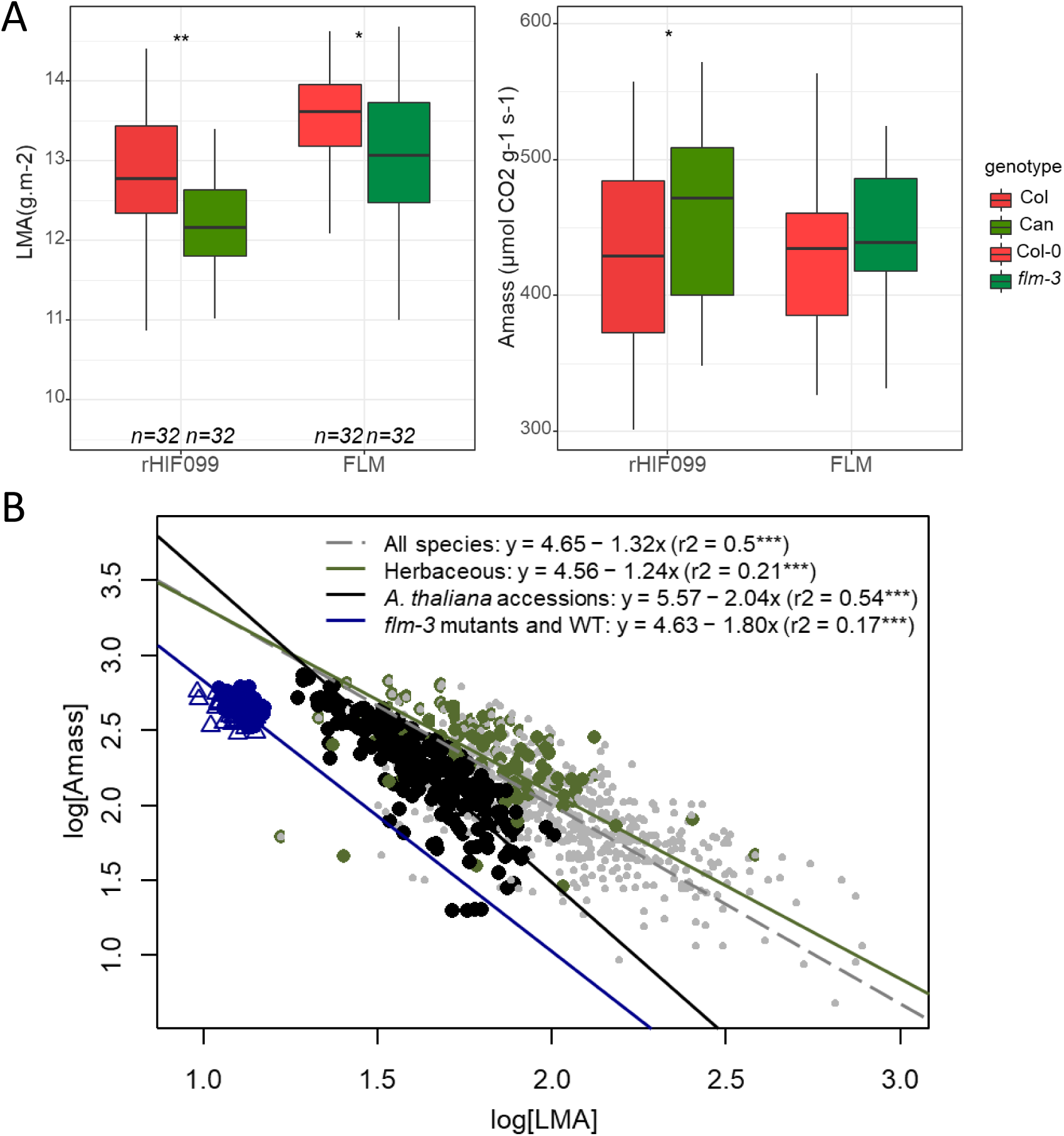
*FLM* allelic variation is modulating important traits of the leaf economics spectrum. A. Boxplots representing the difference of photosynthetic rate by unit of dry mass (*A*_mass_, nmol CO_2_ g^-1^ s^-1^) and leaf dry mass per area (LMA, g m^-2^) between the two allelic lines of rHIF099 or between *flm-3* and Col-0. B. Variations in leaf physiology triggered by *FLM* variation compared to the worldwide leaf economics spectrum, i.e. the relationship between log_10_-transformed *A*_mass_ and LMA. Grey points represent all plant species from the GLOPNET dataset (Wright et al., Nature 2004) (*n* = 2206). Green points represent only herbaceous species from the GLOPNET dataset (*n* = 255). Black circles represent natural accessions of *A. thaliana* (Sartori et al., BioRxiv 2018) (*n* = 378). Blue circles and triangles represent *flm-3* mutants and wild-type measured in this study (*n* = 128). Regressions have been estimated with standard major axis (SMA). *r*^2^: coefficients of correlation from SMA regressions. ***: *P* < 0.001.

The negative correlation between LMA and *A*_mass_ can be attributed to the pleiotropic effect of the *flm-3* mutation on the two traits (Fig. 4A). Moreover, the *FLM* effect on LMA did not depend on the genetic background, as both *flm-3* and rHIF099-Can were significantly different from their respective Col background (Fig. 4A). By contrast, the increase in *A*_mass_ was only significant in the rHIF lines even though a trend was observed in the same direction in the *flm-3* mutant / WT comparison (Fig. 4A). Together, these results suggest that a single mutation at *FLM* has substantial pleiotropic effects on plant physiology. For instance, *flm-3* triggered 2.4% and 2% of the variation observed across all plant species in LMA and *A*_mass_, respectively; and *FLM-Can* triggered 3.5% and 2.2% of the variation observed across all plant species in LMA and *A*_mass_, respectively.

### *FLM* variation relevance in nature

To understand the natural implications of the polymorphism identified, we screened all *A. thaliana* strains with data available for SNP28958 (https://1001genomes.org). Surprisingly, out of the available 50 strains carrying this SNP, 45 come from the Iberian Peninsula (hereafter called IP strains), 2 from France, 2 from Italy and Can-0 from the Canary Islands (Supplemental Table1). To evaluate the impact of SNP28958 on *FLM* expression in natural IP strains, two groups of 9 IP strains carrying either the Can or the Col allele were selected for qRT-PCR experiments. A clear association between the allelic form and the accumulation level of *FLM-β* was observed (Fig. 5A). Moreover, a shift in *FLM-δ* size was only detected in the Can-like IP accessions indicating that this SNP retain the same hypofunctional molecular function as the FLM-Can allele (Fig. S9).

**Figure 5:**
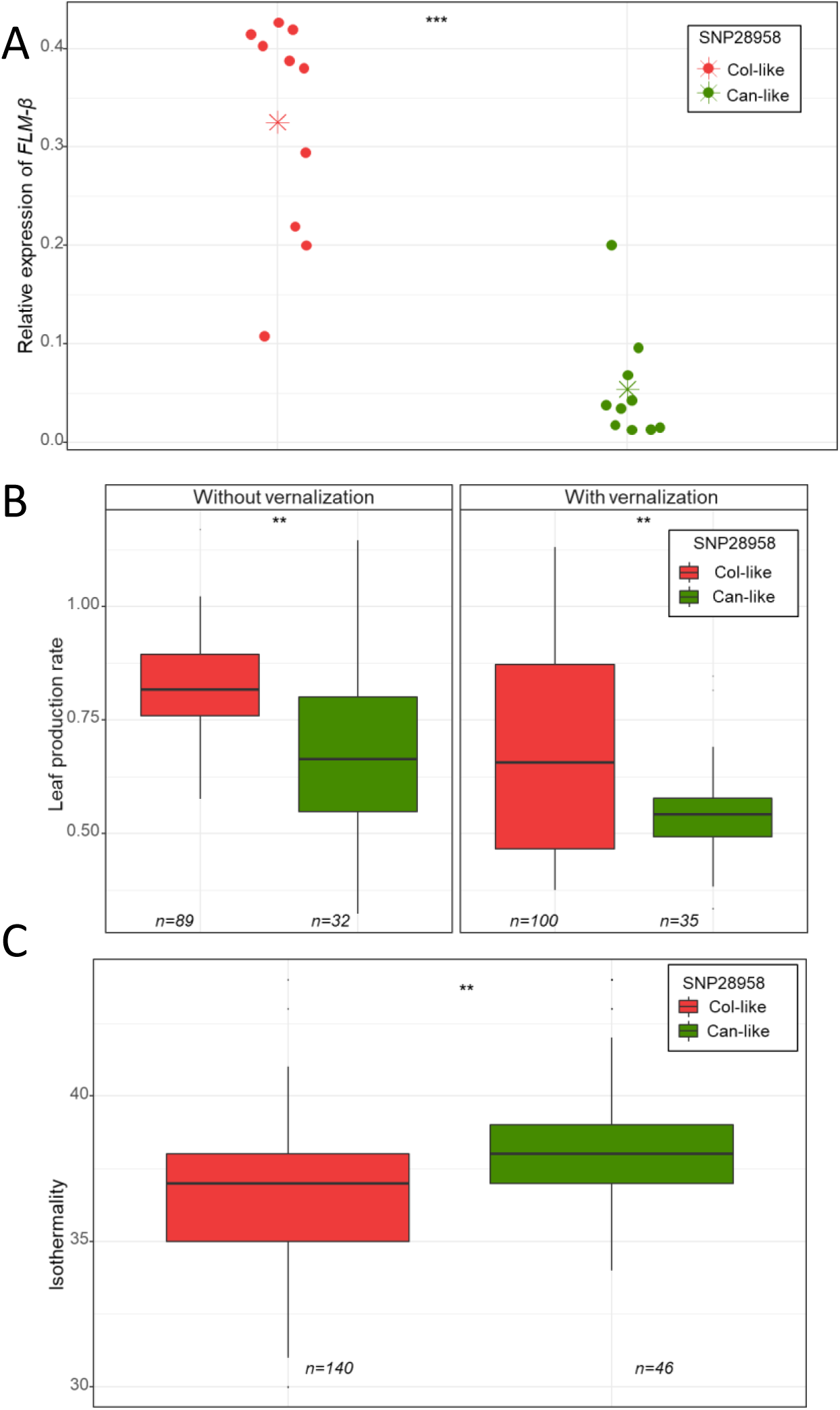
The polymorphism at the position 28,958,437bp in Can-0 modulates *FLM* function and may play a role in the Arabidopsis IP strains adaptation. A- Expression of *FLM-β* was measured by qRT-PCR using the primer pair F3+R3b in 9 Arabidopsis strains from the Iberian Peninsula carrying either the *Can* (green) or the *Col* (red) genotype at the position 28,958,437bp. Each data point represents 1 IP strain coming from a bulk of 12 days old plantlets grown in vitro. The red and green asterisks is the mean expression of each genotype. B- Boxplots represent leaf production rate with or without vernalization in Iberian Peninsula strains categorized according to their genotype at the position 28,958,437bp. The data were obtained from a previous publication (Mendez-vigo et al 2011). Significance of the differences were calculated according to a mixed-effect model using the genotype as a random factor and the kinship matrix as covariates (SupTable3). C- Boxplots represent isothermality in IP strains categorized according to their genotype at the position 28,958,437bp. Isothermality is calculated as follows: Mean of monthly (max temp - min temp) / (Max Temperature of Warmest Month-Min Temperature of Coldest Month) *100, and was obtained from http://www.worldclim.org/. Significance of the differences were calculated according to a mixed-effect model using the genotype as a random factor and the kinship matrix as covariates (SupTable3).

We then wanted to know if the functional impact of SNP28958 on multiple traits could play a substantial role in IP strains adaptation to their habitat. We used a set of 186 IP strains for which we were able to gather geographic and phenotypic data from previous publications including the 46 strains carrying SNP28958 (Mendez-Vigo et al., 2011; Vidigal et al., 2016) (Supplemental Table2). We used this dataset to fit a general linear model corrected by a kinship matrix to avoid bias due to population structure (Supplemental Table3). Consistent with our previous observations, we found significant associations between the *FLM* causal polymorphism and leaf production rate with or without vernalization (Fig. 5B).

Can-like IP strains do not cluster geographically and are scattered across Spain and Portugal (Fig. S10). We used climatic variables (http://www.worldclim.org/) collected over 30 years to know if the location of the Can-like IP strains was associated with particular environmental parameters. We found a significant association between the causal polymorphism and isothermality (p=0.0017), which corresponds to the mean of the monthly temperature range during the day, divided by the maximal variation of temperature over the year (Fig. 5C). This difference seems to be driven mainly by the monthly mean diurnal range for which a similar tendency is observed (p=0.052). This means that the IP strains carrying the *FLM* Can-like allele are rather located in areas where diurnal monthly temperatures are less homogeneous and could be interpreted as a specific adaptation.

### *FLM* allele evolution

Pairwise genetic distances calculation among 1,135 sequenced strains of *A. thaliana* recently showed that 26 of them, referred to as “relicts”, were highly divergent and are supposed to be old lineages coming from ice-age refugia (Alonso-Blanco et al., 2016). Among them, 21 strains form a group coming from the Iberian Peninsula including 13 carrying the SNP28958 (Supplemental Table2). This allele is thus enriched in the relict group (13/21 = 62%) compared to the overall Iberian Peninsula (46/187 = 24.5%; z-test : p=0.00083). A complex pattern of genomic introgressions has recently been highlighted between relict and non-relicts IP strains showing that their genome is rather a mosaic of the 2 groups (Lee et al., 2017). We used this published dataset to estimate the probability of being relict around *FLM* in the IP strains according to the causal polymorphism. Interestingly, overall-relict IP accessions diverge in their probability to be relict specifically around *FLM* according to SNP28958 (Fig. 6). Conversely, non-relict IP strains that are Can-like at *FLM* show a specific increase of probability to be relict at *FLM* position (Fig. 6). It is also noteworthy to mention that Can-0 belongs to an independent relict group (Alonso-Blanco et al., 2016). Furthermore, the FLM-Can allele is also found in 11 of the 14 Madeiran lines that are even more divergent than the relict groups identified in the first place, likely because they have less chance of admixture with non-relict strains (Fulgione et al., 2018). Altogether, these data strongly suggest that the *FLM* allele from Can-0 has an ancestral origin.

**Figure 6:**
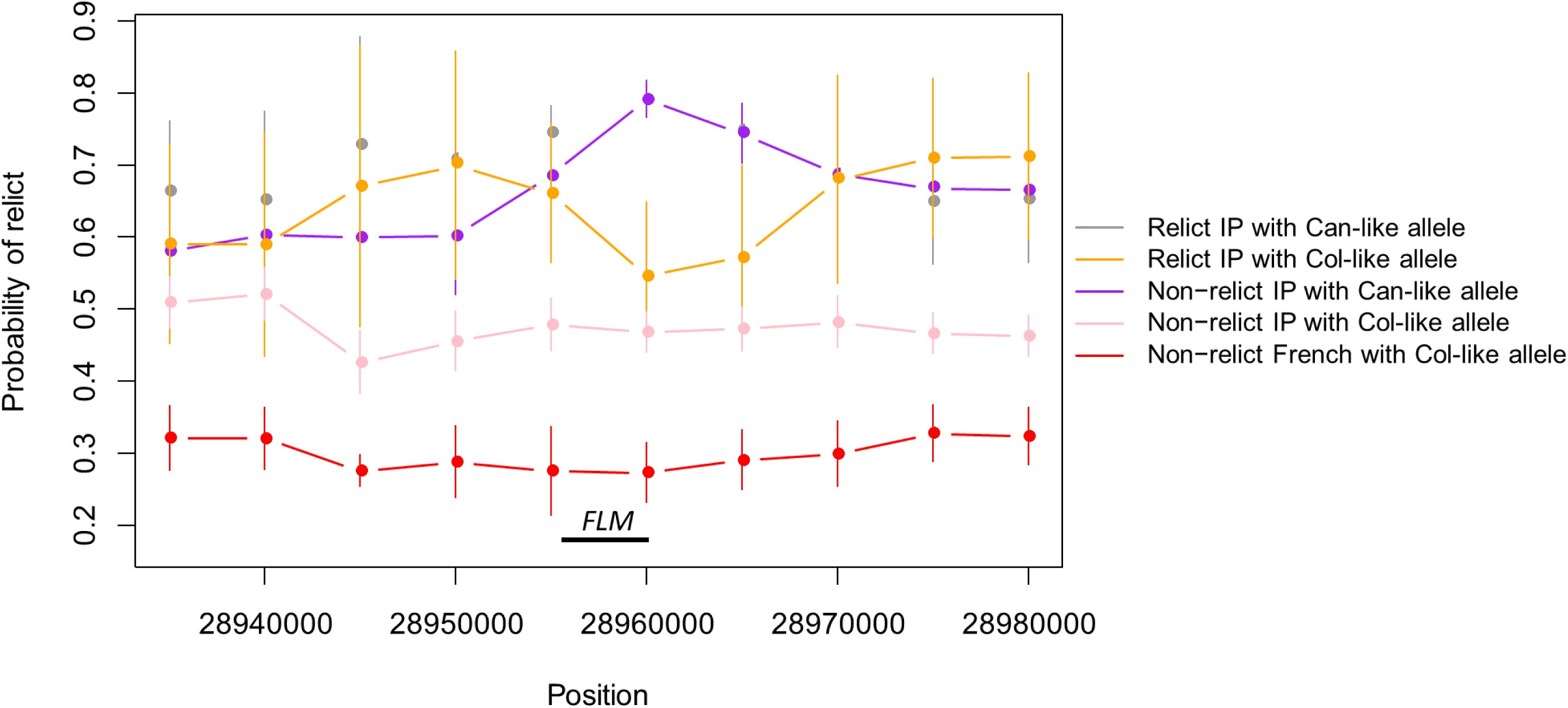
The polymorphism at the position 28,958,437bp is associated with a relict origin. Curves show the average probability of haplotypes to be iberian relicts around *FLM* using an analysis performed previously (Lee et al 2017 Nature Com). Each data point represents the mean over 10kb sliding windows of each category sorted according to their relict origin (relict IP or non-relict IP) and their genotype at position 28,958,437bp (Col-like or Can-like allele). Error bars represent confidence interval at 95%.

## DISCUSSION

Plant growth is a complex trait integrating variation from underlying features called functional traits (Violle et al., 2007) such as net photosynthetic rate, nitrogen metabolism or water use efficiency, themselves also affected by environmental cues throughout the plant life cycle. Plant growth variability is thus the output of the joint action of multiple genes at different levels of organization and is of potential adaptive significance. Likewise, studying this trait is of interest to better understand the genetic architecture and the degree of pleiotropy of traits under complex selective pressures. Here, we intended to map the genetic bases of multiple non-destructive traits related to rosette growth in a RIL population using our phenotyping platform and focused on QTLs co-localized at the end of chromosome 1 having mild effects on growth. We found that this is the result of pleiotropic functions of the *FLM* gene (Fig. 1 & Fig. S1). Further phenotyping experiments showed that *FLM* is also controlling leaf temperature, photosynthetic rate per unit of leaf dry mass and leaf mass per area (Fig. S8, Fig. 4A). Altogether, these results highlight that variation at *FLM* plays a much wider role than initially described.

FLM is a well-known transcription factor whose splicing variation in response to temperature change modulates flowering time (Scortecci et al., 2001; Balasubramanian et al., 2006). *FLM-β* is the repressive variant of this gene, and its down-regulation by increasing temperature triggers flowering (Posé et al., 2013; Lutz et al., 2015, 2017; Capovilla et al., 2017). The *FLM* coding sequences of Col-0 and Can-0 do not differ but *FLM-β* expression is strongly reduced in Can-0 compared to Col-0 (Fig. S3B & Fig. 2B). We identified a single nucleotide substitution in the 2nd intron of this gene (‘SNP28958’), which is creating a seemingly ‘hot’ acceptor splicing site (Fig. S5) and consequently reduces *FLM-β* accumulation in Can-0 as well as in other related natural strains (Fig. 2B & 5A). This leads mainly to the production of transcripts containing a premature stop codon in the 3rd exon (the 2nd exon used for the *FLM-β* variant being skipped). These aberrant variants may then be targeted by nonsense-mediated mRNA decay, as previously reported as a mechanisms repressing *FLM* activity under higher temperature (Kalyna et al., 2012; Sureshkumar et al., 2016). We have specifically validated the impact of the SNP28958 by testing its ability to change the functionality of *FLM-Col* in complementing the *flm-3* mutant phenotype (Fig. S6A).

Intronic polymorphisms are already shown to modulate *FLM-β* expression level in *Arabidopsis* natural strains and thus correlate with flowering time (Lutz et al., 2015, 2017). This gene seems preferentially targeted in non-coding region suggesting that a total loss of function might be too deleterious for the plant. Being able to respond to subtle changes in ambient temperature is indeed crucial to set seeds at the appropriate time of the season. Although very lowly expressed (Fig. 2B), *FLM-β-Can* expression is still responsive to ambient temperature increase (Fig. S4) and our complementation assay shows that the genomic fragment of *FLM-Can* is weakly functional regarding flowering time but also growth (Fig. S6A). Moreover, as we showed that *FLM* controls other traits than flowering time during vegetative growth, its complete inactivation may be maladaptive. Having an *FLM* knock-down (by the substitution SNP28958) instead of a knock-out would allow to keep a certain thermosensitivity while coordinately shifting several other traits important for leaf growth. Loss-of-function mutations were proposed to play a substantial role in plant adaptation (Monroe et al., 2018; Xu et al., 2019). However, the selective value of such natural mutations was rarely demonstrated so far (Poormohammad Kiani et al., 2012; Gujas et al., 2012; Wu et al., 2017). Other work showed that duplication followed by neofunctionalization may also be a source of adaptation (Prasad et al., 2012). Even if it is challenging, this illustrates the value to identify causal genes as well as the causal polymorphisms, as this type of subtle genetic variants are more difficult to reveal using large-scale approaches.

Many examples in human showed how natural variation in alternative splicing is altering gene functions, triggering numerous diseases (Park et al., 2018). In plants, examples are more scarce, yet alternative splicing is occurring in at least 30% of intron-containing genes in several plant species and is likely to play a role in adaptation (Zhang et al., 2010; Reddy et al., 2013; Shen et al., 2014; Thatcher et al., 2014; Mandadi and Scholthof, 2015; Chen et al., 2018). For instance, sequence variation in introns of the PYRROLINE-5-CARBOXYLATE SYNTHASE 1 gene was shown to disturb its proper splicing and consequently modifies proline accumulation, an important trait for drought and freezing tolerance (Kesari et al., 2012). In another example, a single polymorphism in *FLC* intron is changing splicing of the antisense transcript *COOLAIR* and then modulates *FLC* expression (Li et al., 2015). In both cases, an association between functional genetic variation and climatic variables was observed. Our work is contributing to this nascent topic by highlighting the importance of the modulation of gene function through natural variation in non-coding region. Indeed, we found that SNP28958 segregates among *Arabidopsis* strains from the Iberian Peninsula (Supplemental Table2). The categorisation of these strains according to the genotype at this polymorphism allowed to test association with phenotypic and climatic variables. It is noteworthy that strains carrying SNP28958, i.e. the weak allele of *FLM*, which might thus be less responsive to ambient temperature change, are overall located in environments where the amplitude in temperature variation is more pronounced. Therefore, it is conceivable that this *FLM* allele is advantageous to protect plants against fluctuating temperature and avoid the inappropriate triggering of flowering. Genetic variation affecting thermosensitivity of splicing in a key clock gene was proposed to drive thermal adaptation from equatorial to temperate climate in *Drosophila* species, supporting a similar hypothesis (Low et al., 2008).

Adaptation to fluctuating thermal environments could also be associated with faster growth in order to complete the life cycle before stress occurs (Grime, 1977). Consistently, further phenotypic inspection showed that *FLM* also influences functional traits reflecting fundamental aspects of leaf physiology, plant growth and reproduction (Wright, 2004; Reich, 2014; Díaz et al., 2016). Most notably, *FLM* modulated photosynthetic rate and leaf dry mass per area (Fig. 4A), two core traits of the universal trade-off between carbon acquisition and nutrient retention, called the leaf economics spectrum (LES). The fact that our absolute values for these traits fell outside of the range previously observed across species and among *A. thaliana* accessions can be explained by the growth conditions in which plants from the *flm* dataset were measured (Fig. 4B). Indeed, plants were grown in the greenhouse under relatively low light intensity (*ca*. 65 *µ*mol m^-2^ s^-1^ PPFD), compared to the natural accessions grown in the PHENOPSIS phenotyping platform under moderate light intensity (*ca*. 195 *µ*mol m^-2^ s^-1^ PPFD; Granier et al., 2006), or the interspecific data collected in common garden experiments, presumably under much higher (and fluctuating) light intensity. Still, the relationship between these traits was conserved in the *flm-3*/Col-0 comparison (*p*>0.05, *r*^2^=0.17). The loss of function of *FLM* triggered coordinated changes in functional traits that incrementally re-oriented whole-plant physiology toward the resource-acquisitive side of the LES. Strikingly, the impact of this *FLM* mutation in *A. thaliana* may contribute to 2-3% of the worldwide variation in plant physiology along the LES (Wright et al., 2004). *FRIGIDA* has been documented in plants as a gene with comparable level of pleiotropy on physiology and life-history (Lovell et al., 2013). *ERECTA* had also been associated with pleiotropic functions relating reproductive development and plant physiology (Masle et al., 2005). However, the phenotypic contribution of these genes was not assessed in the context of the LES.

Two different scenarios may explain the pleiotropy exerted by *FLM*. On the one hand, as physiological traits are highly interconnected (Angert et al., 2007; Lovell et al., 2013), *FLM* would modulate directly one of them e.g. water fluxes in the plant, which in turn would indirectly change the C requirements, the photosynthetic rate and finally plant growth. On the other hand, *FLM* may activate or repress a set of target genes, selected through evolution owing to physiological constraints along the spectrum of leaf economics. Investigating the transcriptional network controlled by *FLM* might provide some answers as previously performed for *FLC* (Deng et al., 2011). In parallel, it may be worth looking at the transcriptome according to *FLM-β* expression level. Another unanswered question lies with the role of *FLM-δ* which is expressed at a substantial level in many *Arabidopsis* strains but was not shown to play an essential role in modulating flowering time (Lutz et al., 2015, 2017; Capovilla et al., 2017).

According to the pleiotropic role of *FLM* on functional traits such as photosynthetic capacity or LMA, placing the *Arabidopsis* strains carrying SNP28958 toward a rather more resource-acquisitive side of the LES, it is tempting to speculate that the associations between the *FLM* genotype and either growth strategy (leaf production rate) or the environment (isothermality) is of adaptive significance. Interestingly, the knock-out (*flm-3*) and the knock-down (Can) alleles of *FLM* used here display many features of the drought escape strategy, one of the three strategies developed by the plants to overcome water shortage (Ludlow, 1989). Indeed, these genotypes have a more rapid vegetative development, earlier flowering and higher photosynthetic rate, which would help them to complete their life cycle before a severe drought season (Kooyers, 2015). Weak genotypes at *FLM* have also cooler leaf temperature indicating that WUE may be lower in these plants. This characteristic would be advantageous in environments where water is either not limiting or discontinuous, either to ensure non-limiting water acquisition or to acquire it faster, respectively. A recent genome-environment association study performed on local populations of *Arabidopsis* growing in south west of France identified *FLM* as a potential candidate associated with the amount of precipitation (Frachon et al., 2018). We did not observe associations with precipitation variables but our dataset differs from Frachon *et al.* (2018) as it covers the whole Iberian Peninsula potentially encompassing a much wider range of genetic and environmental variation. Evaluation of the fitness consequences driven by natural allelic variation in the gene *MITOGEN ACTIVATED PROTEIN KINASE 12*, which modulates WUE, showed that low WUE genotype were advantageous in the face of competitors (Campitelli et al., 2016). It would be interesting to investigate the adaptive value of *FLM* by combining water availability, temperature fluctuation and competition using our *Arabidopsis* transgenic lines differing only at SNP28958. Reciprocal transplant experiment could also be considered to quantify the adaptive effect of the *FLM* alleles (Ågren and Schemske, 2012).

The scenario of *Arabidopsis* migrations is regularly revisited as more data are available for population genetic studies (Fulgione and Hancock, 2018; Hsu et al., 2019). In many instances, Iberian Peninsula is consistently mentioned as a location where highly diverged *Arabidopsis* lines (called relicts) are found in preserved and ancestral habitats (Sharbel et al., 2000; Brennan et al., 2014; Alonso-Blanco et al., 2016). We identified SNP28958 in the Can allele of *FLM* and we found it in some IP strains i.e in 2 of the 5 relict groups identified (1001 genome). Interestingly, the frequency of SNP28958 is enriched 2.5 times among the relict IP strains compared to all IP strains. Furthermore, even among the non-relict IP strains carrying SNP28958, the genotype probability to be relict is specifically rising at the *FLM* locus, further supporting its relict origin. Finally, we found SNP28958 in 11 of the 14 Madeiran lines which are considered as some of the most divergent *Arabidopsis* strains so far (Fulgione et al., 2018). *Arabidopsis* migrations were associated with climate disruptions such as temperature rise or precipitation fluctuation (Fulgione and Hancock, 2018). Regarding the pleiotropic functions of SNP28958 towards more acquisitive strategy, it is conceivable that this knock-down allele was adapted to ancestral habitats and was then outcompeted and wiped out by the *FLM-Col* allele during the last expansion of non-relict *Arabidopsis* in Europe (Alonso-Blanco et al., 2016; Lee et al., 2017).

Since Darwin’s studies, the classical view in evolutionary biology is that adaptive evolution occurs through gradual steps and is therefore rather slow and highly polygenic (Darwin 1859; Fisher, 1958). However, an increasing number of studies showed that evolution of organisms can happen through more steep step for examples due to polyploidization events or to mutations in genes of large effect (Sheehan et al., 2016; Van De Peer et al., 2017; Bradshaw Jr and Schemske, 2003). Under strong selective pressure, it seems logical that adaptation must be accelerated through new large effect alleles for simple traits or by targeting pleiotropic genes if the given trait is more complex. While pleiotropy (and its role in adaptation) has been studied mainly theoretically since years, few empirical works relying on published mutant data or QTL analysis showed that pleiotropic genes are relatively rare in a range of organisms (Wagner et al., 2008; Wang et al., 2010). Recent works demonstrated that a combination of intermediate pleiotropic loci having a specific function may constitute an optimal balance for a relatively fast adaptation and/or to manage complex traits, but the functional validation of candidate genes was not yet done (Frachon et al., 2018; Fusari et al., 2017). Our study is in line with those works, and provide an important case study, as natural variation of *FLM*-coordinated functions is associated with contrasted life strategy along the LES axis and is associated with the environment. Such a coordinated pleiotropy caused by a single gene drives a whole module of traits and might be more responsive in the face of climatic challenges.

## MATERIALS & METHODS

### Genetic material

A segregating population (named 19RV) derived from the cross between Can-0 (accession 163AV) and Col-0 (accession 186AV) is described and available from the Versailles Arabidopsis stock centre (http://publiclines.versailles.inra.fr/; Simon et al., 2008)). Description and associated data are gathered at http://publiclines.versailles.inra.fr/page/19. 358 RILs were phenotyped in similar experimental design as previously described (Marchadier et al., 2019).

HIF was fixed as previously described (Marchadier et al., 2019; Loudet et al., 2005) from the following RIL (19RV337) with a residual segregating region covering position 27,401,322bp to 29,693,733bp on chromosome 1 (See Supplemental Table4). For fine-mapping, 3,360 progenies of heterozygous 19HV337 were genotypically screened to find recombinant individuals within the candidate region (rHIF). By interrogating the segregation of PRA29 in informative rHIFs, the candidate interval for the QTL was reduced to approximately 35 kb. The detailed genotype of 19HV337 and its rHIFs can be found in Supplemental Table4.

The mutant *flm-3* is a SALK T-DNA line kindly provided by Markus Schmid (Stock Centre number N641971) Quantitative complementation assay was performed as described previously (Mackay, 2004; Loudet et al., 2007; Vlad et al., 2010) by crossing each rHIF allelic line to either the flm-3 mutant or its WT background (Col-0) and testing its phenotype directly in the F1 plants.

### Phenotyping and growth conditions

Initial RILs phenotyping, fine-mapping and mutant characterization were first obtained on the Phenoscope robots (Tisné et al., 2013) in well-watered conditions (60% soil water content saturation) as previously described (Marchadier et al., 2019). The RIL set and the parental accessions have been phenotyped in 2 independent Phenoscope experiments, with 1 individual plant per RIL per condition per experiment. The phenotypic values can be found in the Supplemental Table5. The growth room is set at an 8 hours short-days photoperiod (230 μmol m-2 sec-1) with days at 21°C/65%RH and nights at 17°C/65%RH. Growth-related traits are extracted from daily images after segmentation (PRA = Projected Rosette Area = cumulative growth ; RER = Rosette Expansion Rate = relative growth rate), as well as parameters describing the rosette colour expressed in the HSV (Hue/Saturation/Value) scale.

The growth chambers used for the experiments carried out at 16°C and 23°C was set at a 16 hours long days photoperiod 150 μmol m-2 sec-1, 65%RH.

For the in vitro culture, seeds were surface-sterilized for 10 minutes in 70% EtOH, 0.1% TritonX-100, followed by one wash with 95% EtOH for another 10 minutes. Sterile seeds were then resuspended in a 0.1% agar solution and stratified in the dark at 4°C for 3 days. Ten seeds per square Petri dishes (120 mm) containing typical Arabidopsis media (Trontin et al., 2014) were placed on one side and the Petri dishes set vertically. Plants were grown for 12 days in a culture room (21°C, 16 hours light/8 hours dark cycle).

### QTL mapping

QTL mapping was performed using Multiple QTL Mapping algorithm (MQM) implemented in the R/qtl package (Broman et al., 2003; Arends et al., 2010). At first, genotypic missing data were augmented. Then, one marker every three markers were selected as cofactors and significant ones were selected through backward elimination (backward selection of cofactors). QTL was moved along the genome using these pre-selected markers as cofactors, except for the markers in the 25.0 cM window around the region of interest. QTL were identified based on the most informative model through maximum likelihood. According to permutation tests results, a LOD threshold of 2.4 was applied to identify significant QTL.

### Measurement of physiological traits in greenhouse

*flm-3* knock-out mutant and its WT (Col-0), as well as one recombinant HIF segregating for *FLM* (rHIF099) were grown in greenhouse at the Centre d’Ecologie Fonctionnelle et Evolutive (CEFE, Montpellier, France) in fall 2017. Plants were grown in 32 replicates per genotype. Seeds were sown in 260 ml individual pots filled with a 1:1 (v:v) mixture of loamy soil and organic compost, and stratified at 4 °C for three to ten days. At the emergence of the first two true leaves, plants were thinned to keep only one plant per pot. Pots were randomly distributed among four trays that were rotated every day in the greenhouse. All pots were watered twice a week. To reduce environmental heterogeneity in the greenhouse, walls were painted in white and a semi-transparent curtain was installed below the glass roof. Additional light was provided to reach ca. 65 *µ*mol m^-2^ s^-1^ PPFD. Photoperiod and temperature were kept constant at 12 h day length, and 18/16 °C day/night, respectively.

Gas exchanges were determined 28 days after the end of stratification, that is, just before harvest. The rate of CO_2_ assimilation per unit dry mass (*A*_mass_, nmol CO_2_ g^-1^ s^-1^) was measured using a whole-plant chamber designed for Arabidopsis (Li-Cor 6400-17, Li-Cor Inc., Lincoln, NE, USA) connected to a gas analyzer system (LI-6400XT; Li-Cor). *A*_mass_ was determined at steady state at 180 *µ*mol m^-2^ s^-1^ PPFD, 20°C) and at 390 ppm reference CO_2_. After gas exchange measurement, rosettes were cut, leaves were separated and scanned individually for measurements of leaf area (ImageJ 1.43C). Leaf blades were then oven-dried at 65°C for 72 h, and their dry mass (DM) was determined. Leaf dry mass per area (LMA, g m^-2^) was calculated as total blade DM divided by total blade area. The raw data can be found in the Supplemental Table6.

### Cloning procedures

*FLM* genomic fragments of Col-0 and Can-0 were amplified using Phusion high-fidelity Taq polymerase (Finnzymes, http://www.thermoscientificbio.com/finnzymes/) with a couple of primers including Gateway sites and flanking positions -2219 to +4737 relative to the *FLM* start codon of Col-0 annotated in TAIR10 (Supplemental Table7). The pDONR207 entry vector containing the gFLM from Col-0 was mutagenized by PCR to substitute the nucleotide at the position +2759 by a “A”. Gateway-compatible fragments were cloned into the pDONR207 entry vector (Invitrogen) via BP recombination and subsequently transferred into the binary vector pFAST via a LR reaction following the Gateway cloning procedure (Invitrogen, www.invitrogen.com ; Schlamenbach et al 2014, BMC Plant Biology). After a verification done by Sanger sequencing, expression constructs were transformed into Agrobacterium tumefaciens strain C58C1. *flm-3* mutants were then transformed by floral dipping (Clough and Bent, 1998) and transgenic plants were isolated using seed fluorescence. We selected four independent transgenic lines carrying the Col-0 and the Can-0 fragment of *FLM*, and two independent transgenic lines carrying the mutated version of *FLM*.

FLM transcripts variants of Col-0 and Can-0 were amplified with primers including start and stop codons using Phusion high-fidelity Taq polymerase. PCR products were cloned with the Zero Blunt TOPO PCR cloning kit (invitrogen) following manufacturer specifications. Colony PCR was used to assess the size and the frequency of the different isoforms.

### Gene expression analysis

The RNeasy Plant Mini kit (*Qiagen*) was used for RNA extractions followed by a DNAse treatment (*Fermentas*). RT-PCR was performed on 500 ng of RNA using RevertAid H Minus reverse transcriptase (Fermentas) with oligo(dT) in 20*µ*Lreactions. Then, 5 μl of 10-fold diluted cDNAs was used for qRT-PCRs using a CFX96 real-time PCR machine (*BioRad*) with a SYBR solution (*Eurogentec*) using primers listed in supplemental Table7. Expression levels were normalized against the Arabidopsis *PP2A* gene (*At1g13320*).

### Statistical analyses

Significance of the difference in the box plots were calculated with the Mann-Whitney-Wilcoxon test for the pairwise comparisons (‘.’ *p*<0.1; ‘*’ *p*<0.05; ‘**’ *p*<0.01; ‘***’ *p*<0.001) or with the Tukey HSD test for multiple comparisons, the different letters representing groups at *p* < 0.01. Linear regressions between *A*_mass_ and LMA were examined with standard major axis (SMA), using the package *smatr* in R (Warton *et al.*, 2012). Physiological traits (*i.e.* LMA and *A*_mass_) across Arabidopsis accessions were obtained from Sartori et al., 2018, while physiological traits from interspecific data were obtained from the GLOPNET database (Wright *et al*., 2004). All analyses were performed in R 3.2.3 (Team RC 2014).

## ACKNOWLEDGEMENTS

We thank Yann Serrand for the supervision of the *Phenoscope*, and Lilian Dahuron for expert care of our plants. We also would like to thank Markus Schmid (MPI Developmental Biology, Tübingen, Germany; Umea Plant Science Center, Umea, Sweden) who provided seeds of the *flm-3* mutant; Carlos Alonso-Blanco for sharing his experience in sequence diversity analyses; Cheng-Ruei Lee (National Taiwan University) for sharing his analysis concerning relict accessions.

---

**Supplemental figure 1:**
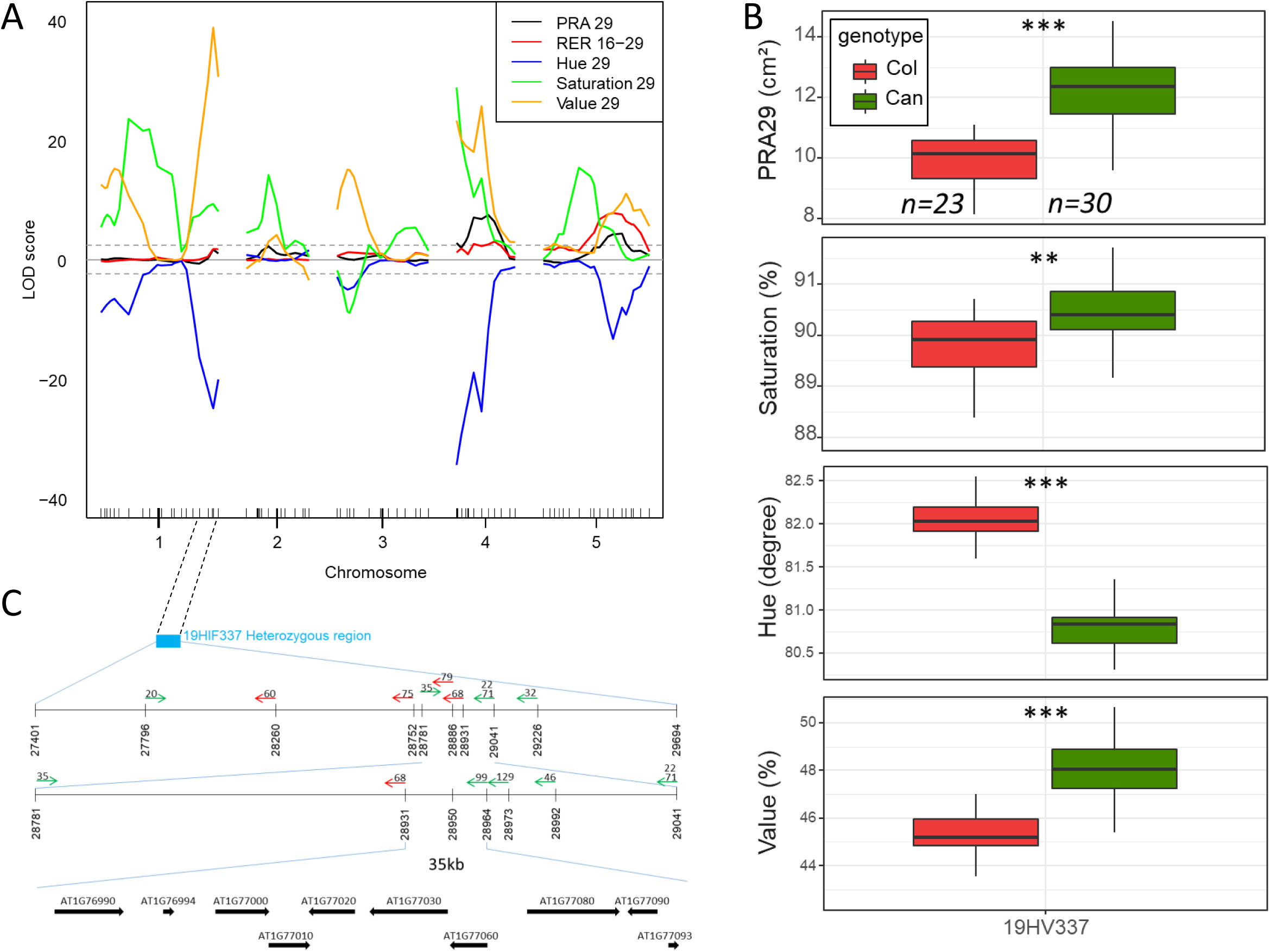
QTL mapping and fine mapping of the QTLs located at the end of chromosome 1. A- QTLs from the Can-0 x Col-0 RIL set for 5 traits: Projected Rosette Area 29 days after sowing (PRA29), Relative Expansion Rate from 16 to 29 days after sowing (RER16-29) and average rosette color expressed in the HSV (Hue, Saturation and Value) scale. Positive or negative LOD score values indicate that the *Can* allele or the *Col* allele increases trait value, respectively. Significance threshold (LOD=2.4) is shown as dotted lines. B- Validation of the QTLs located at the end of chromosome 1 using the heterogeneous inbred family 19HV337 segregating for the region highlighted with the blue rectangle in panel C. The box plots for PRA29, Hue, Saturation and Value show the distribution of the phenotypic data and significance of the genotype effect. C- Analysis of PRA29 segregation in series of recombinants issued from 19HV337[Het] reduced the QTL candidate interval to 35kb. Each arrow is located at the recombination breakpoint of each recombinant line (rHIF). The back of the arrow indicates the fixed (homozygous) region whereas the head points toward the heterozygous region. Green and red colors indicate whether PRA29 is segregating or not in this rHIF, respectively. The thicker arrows at the bottom represent the candidate genes present in the final 35kb-interval of interest.

**Supplemental figure 2:**
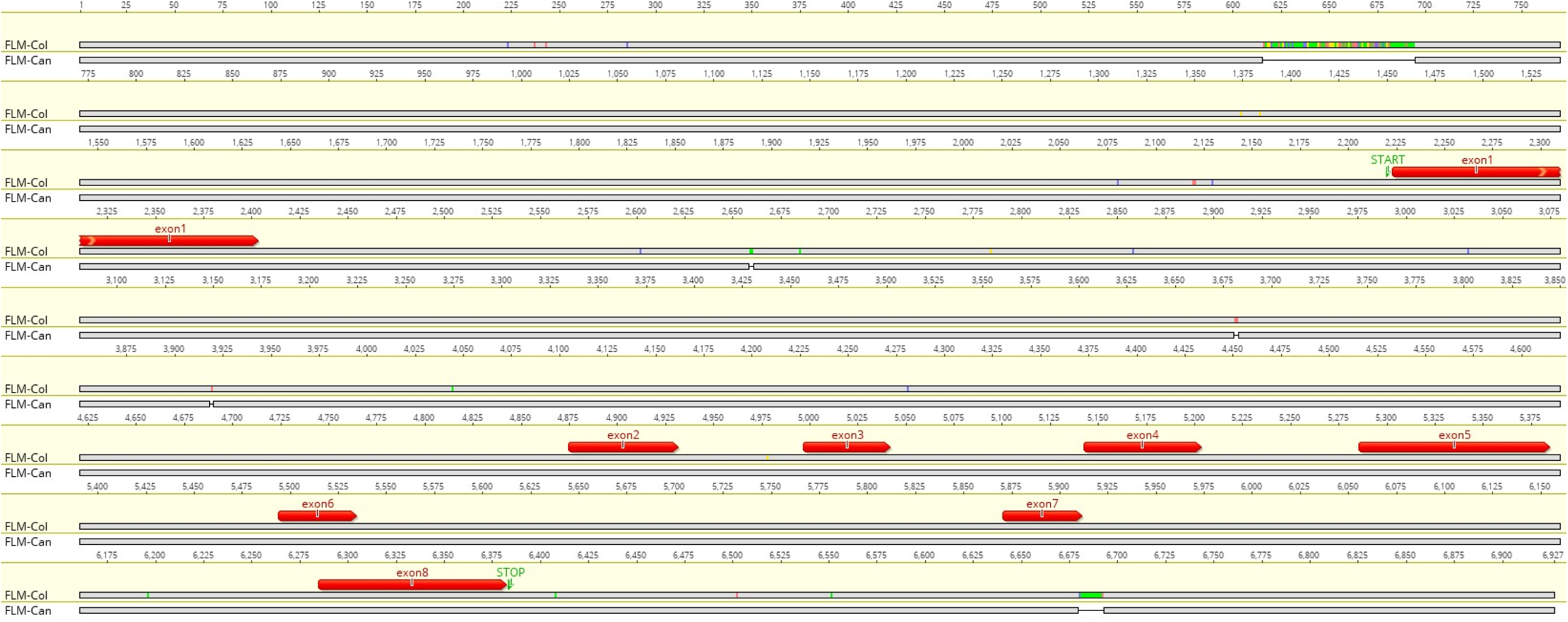
Alignment of the genomic sequences of the *FLM* genes of Col-0 and Can-0. The alignment was performed with the Geneious software. Polymorphisms are highlighted in the top row in which *FLM-Col* is used as a reference. Exons are depicted in red, START and STOP codons in green.

**Supplemental figure 3:**
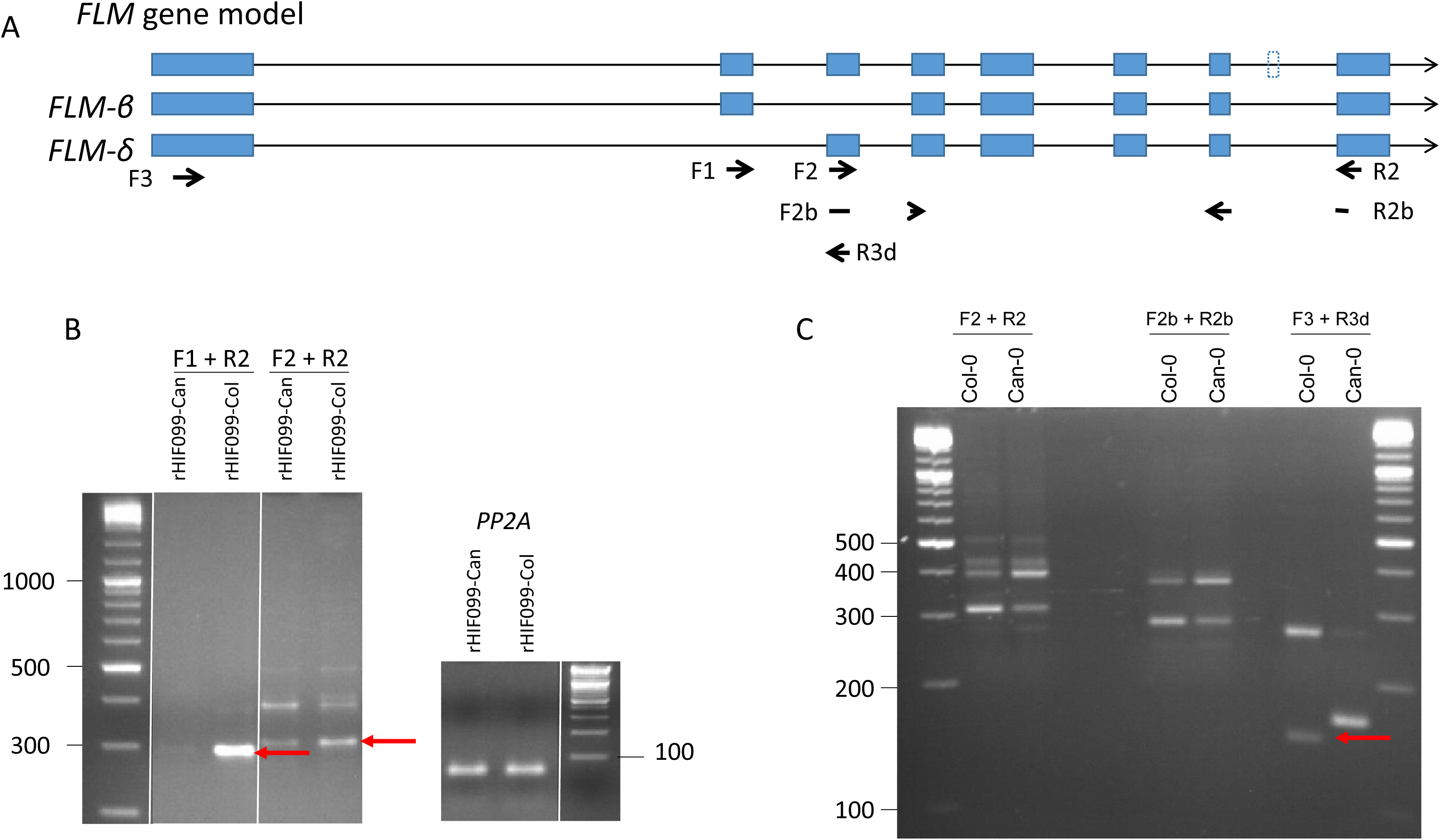
Variation in alternative splicing of *FLM* in Col-0 and Can-0. A. Graphic representation of the *FLM* gene and transcript model as well as its two main splicing variants *FLM-β* and *FLM-δ*. The primers used for semi-quantitative RT-PCR are represented by the arrow. B. Expression level of the *FLM* transcripts characterized by semi-quantitative RT-PCR performed on cDNA obtained from rosette leaves of the rHIF099 homozygous for the *Can* or the *Col* allele. The plants were harvested from a *Phenoscope* experiment 29 days after sowing. Four individuals of each genotype gave similar results as the one presented here. The red arrows indicate the expected size in Col-0. *PP2A* was used as a control gene. The molecular-weight size marker on the side of the gels are expressed in bp. C. Characterization of the *FLM-*□ isoform by PCR carried out on cDNA obtained from rosette leaves of plants grown at 23°C. The red arrows indicate the expected size in Col-0. The molecular-weight size marker on the side of the gels are expressed in bp.

**Supplemental figure 4:**
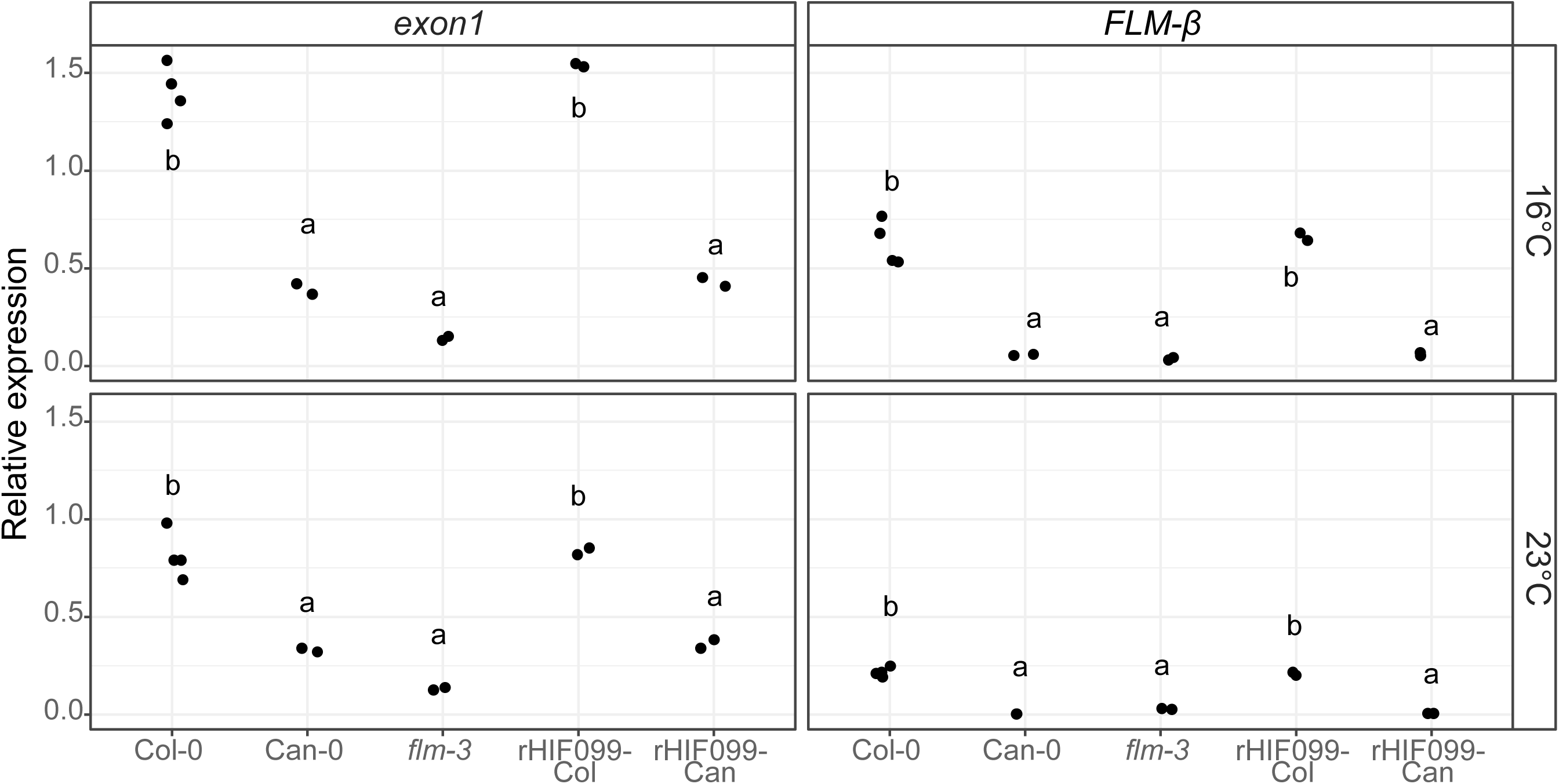
*FLM-Can* accumulation is down-regulated irrespective of the temperature or the genetic background. Relative expressions of the *exon1* and *FLM-β* were measured by qRT-PCR using the primer pairs F3+R9 and F3+R3b respectively (See SupFig3A). Each data point represents cDNAs obtained from pools of 3 rosettes grown at 23°C and 16°C (in long days).

**Supplemental figure 5:**
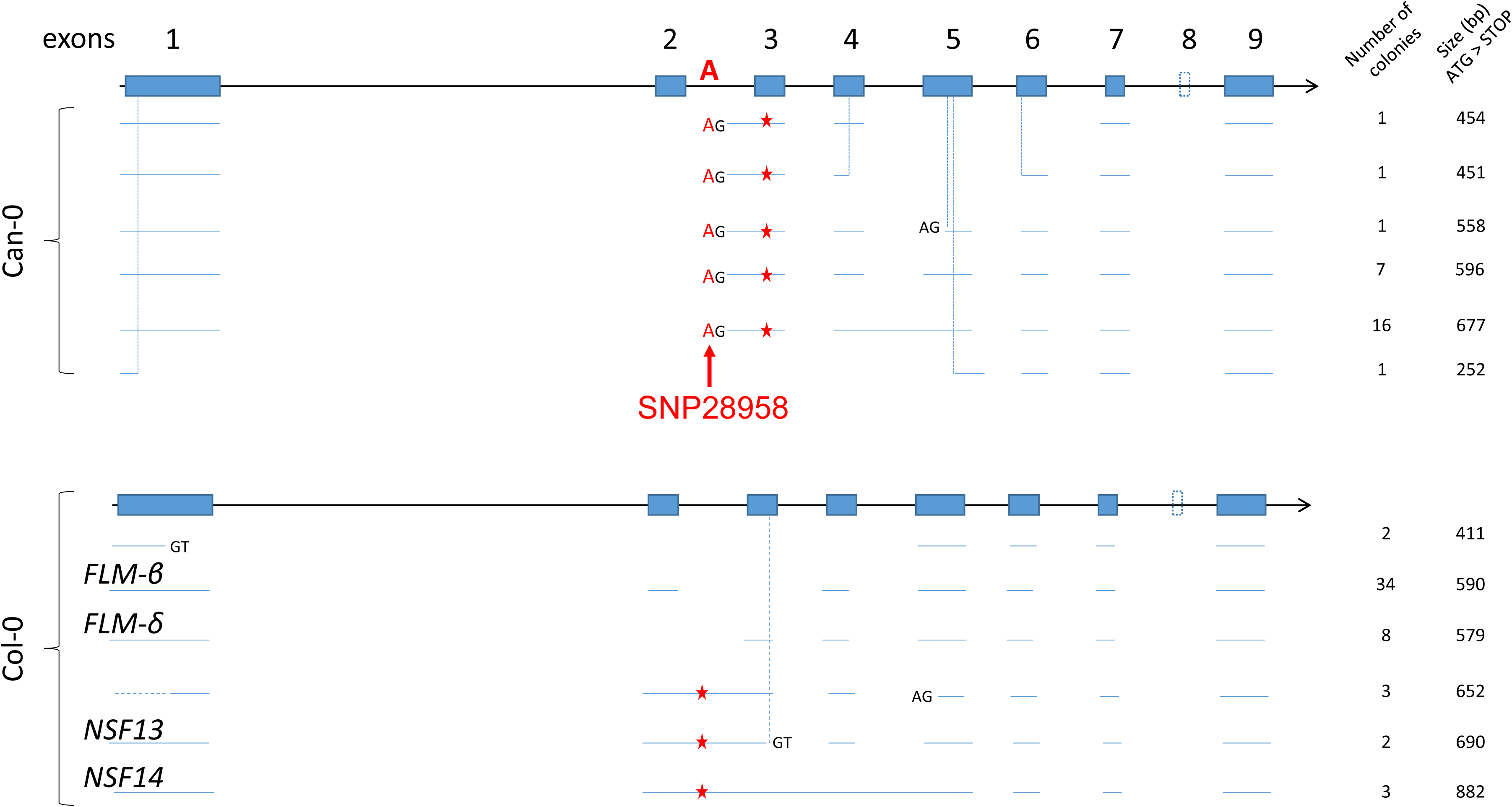
Characterization of the *FLM* isoforms produced at 23°C in Col-0 and Can-0. cDNAs of Col-0 and Can-0 were obtained from pools of 3 rosettes grown at 23°C and were used for the PCR amplification of the *FLM* coding region from the start to the stop codons. Each PCR product was then used for subcloning and the colonies obtained were screened by PCR to estimate the size of many subcloned fragments. Plasmids of one or two colonies carrying a PCR product with a distinct band size were sent for Sanger sequencing. This provided the sequences illustrated here with the horizontal blue lines. An estimation of the quantity of each isoform sequenced is given according to the size of the PCR product obtained during the colony screening. The vertical dashed lines specify a discrepancy between the annotation and the isoform sequenced. New donor (GT) and acceptor (AG) splicing site are depicted when present. Red stars represent premature stop codon. In Col-0, *FLM-β* and *FLM-δ* as well as variants previously identified by Sureshkumar et al 2016 are mentioned (*NSF13* and *NSF14*). The candidate polymorphism responsible for the splicing shift observed in most of the transcripts of *FLM-Can* is depicted in red (‘SNP28958’).

**Supplemental figure 6:**
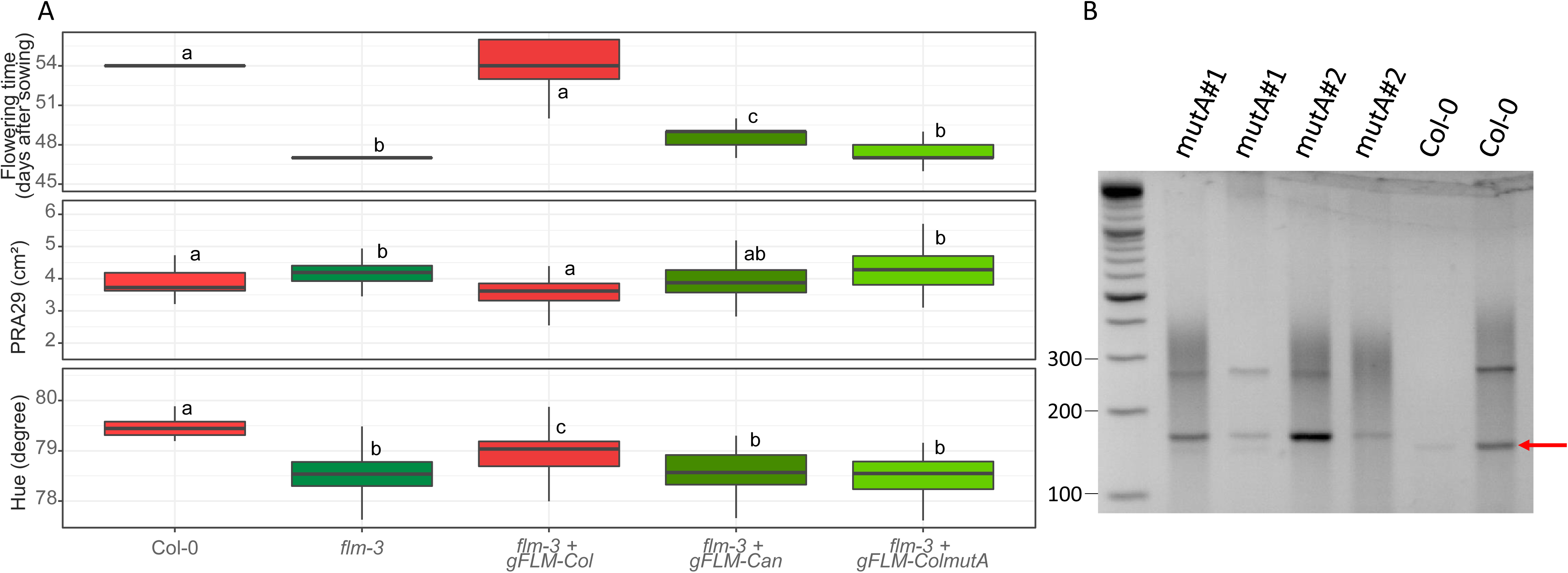
Functional complementation of *flm-3*. A. Different genomic fragments (promoter to 3’UTR) of *FLM* were used to transform the *flm-3* mutant for functional complementation assays: the fragments of Can-0 (gFLM-Can), Col-0 (gFLM-Col) and Col-0 substituted with SNP28958 (gFLM-ColmutA).The data presented here as boxplots for flowering time, PRA29 and Hue were obtained from T2 plants (4 independent lines for gFLM-Can and gFLM-Col, 2 independent lines for gFLM-ColmutA). B. Characterization of the *FLM-*□ isoform by PCR using the primer pair F3+R3d (SupFig3A) in 2 individuals of each gFLM-ColmutA T2 lines and 2 Col-0 plants as a control. The red arrows indicate the expected size in Col-0.The molecular-weight size marker on the side of the gel is expressed in bp.

**Supplemental figure 7:**
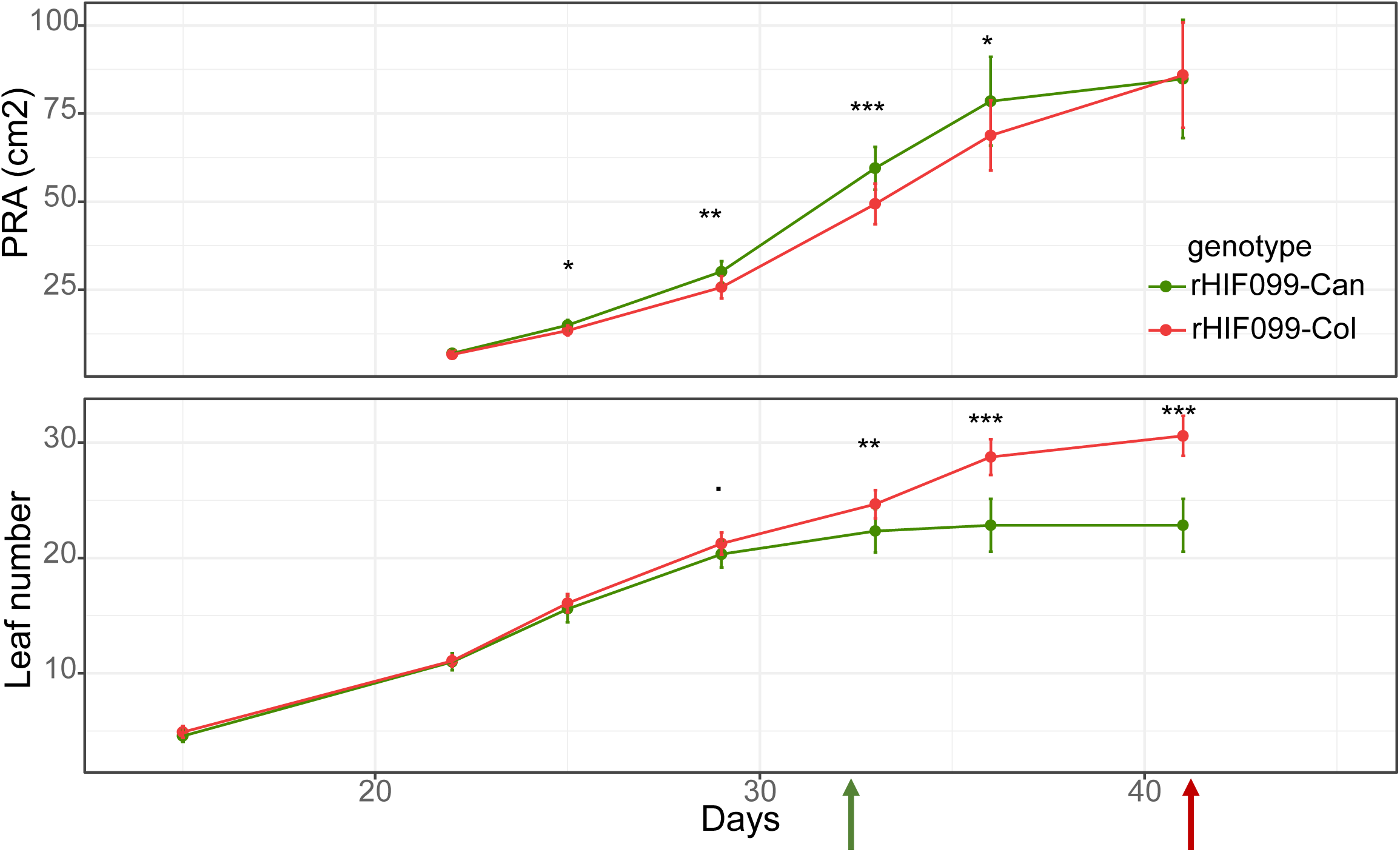
Relationships between vegetative growth and flowering time according to the *FLM* allele. Dynamics of projected rosette area (PRA) and leaf number followed until 41 days after sowing. Green and red arrow indicate the bolting time (floral stem 1cm) in rHIF099-Can and rHIF099-Col respectively.

**Supplemental figure 8:**
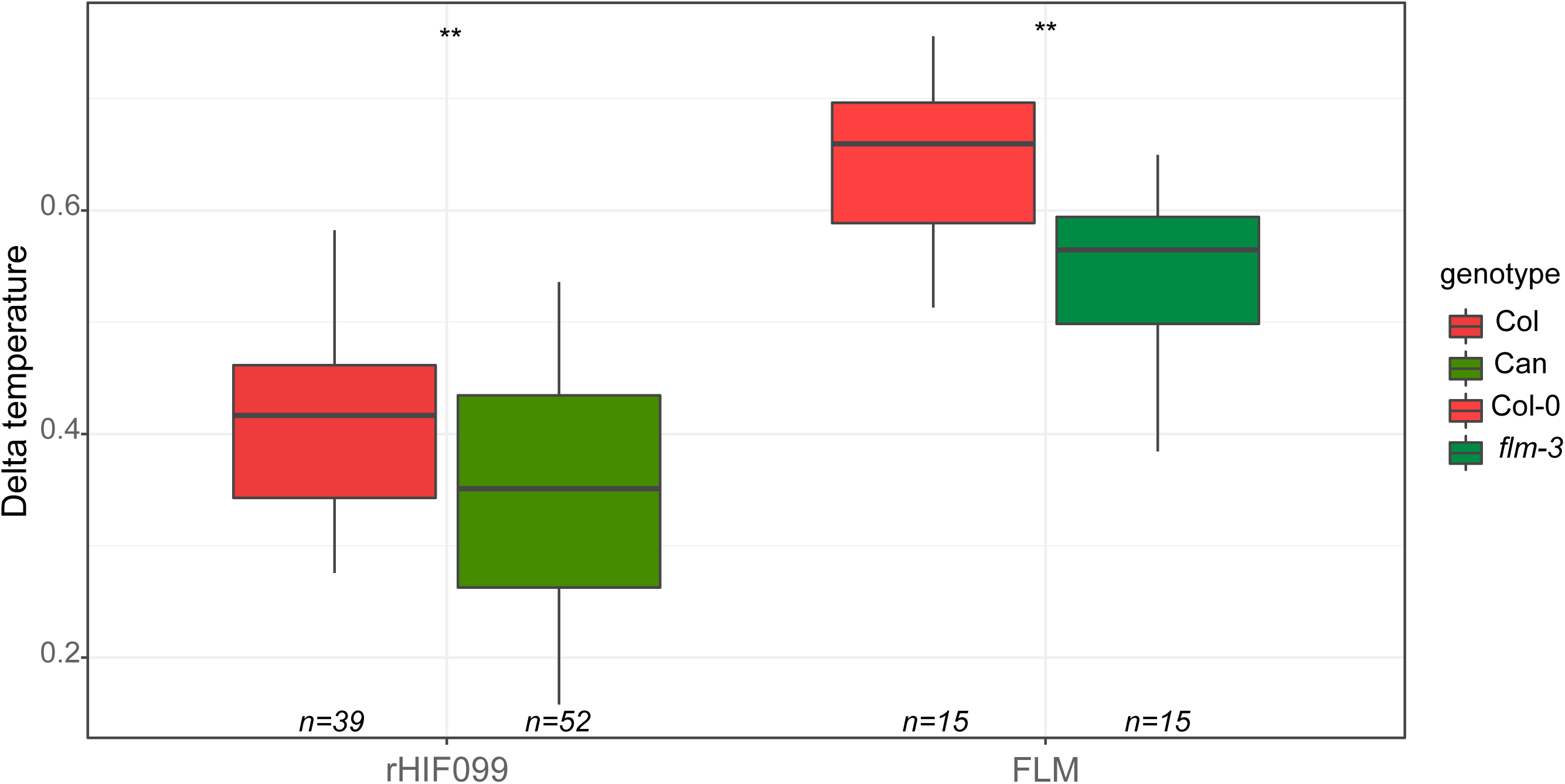
Leaf temperature is affected by the genotype at *FLM*. The data points presented as boxplots were obtained by normalizing the mean of rosette temperatures with the room temperature.

**Supplemental figure 9:**
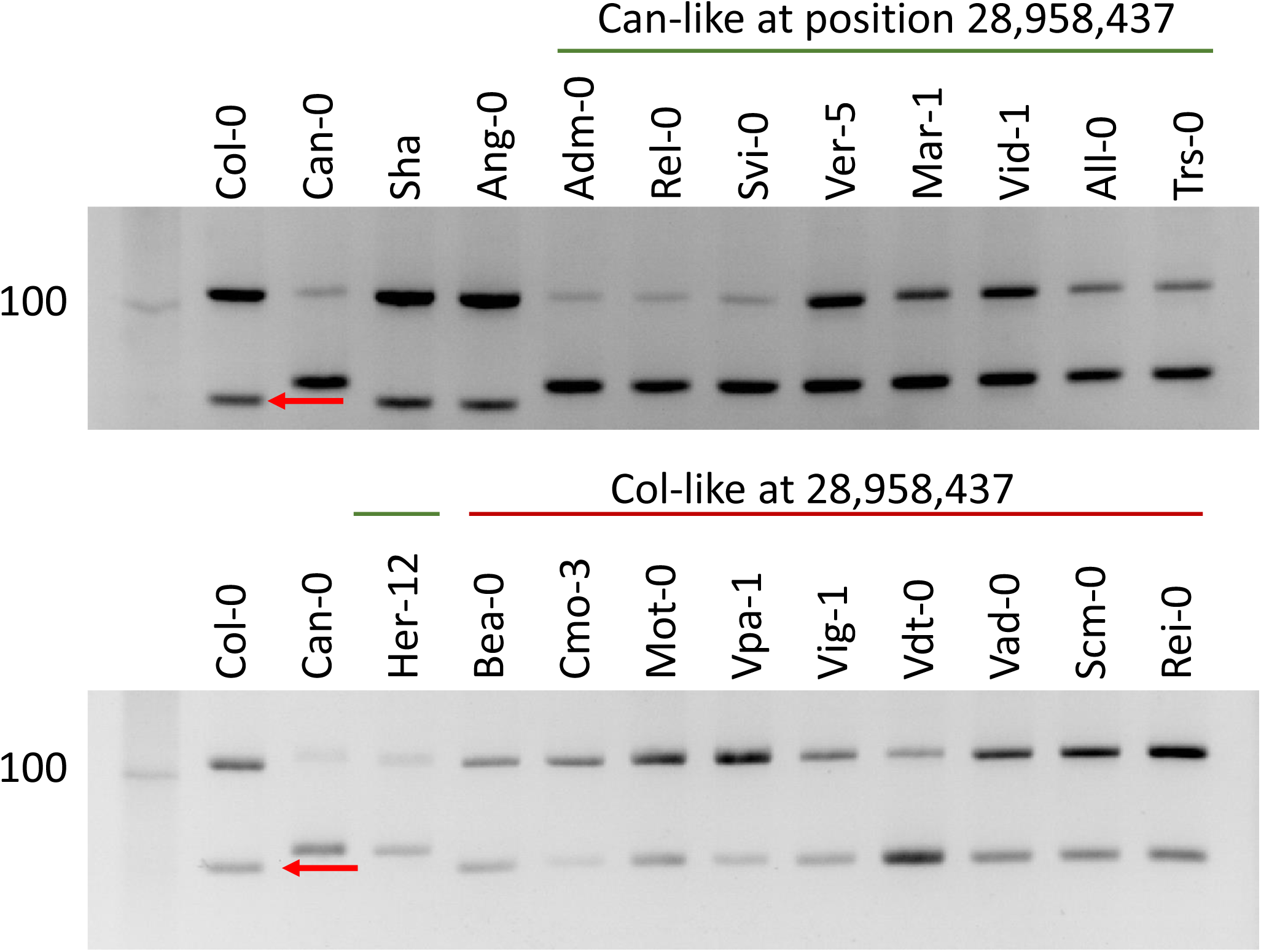
The polymorphism at the position 28,958,437bp changes splicing in natural IP strains. Characterization of the splicing shift by PCR using the primer pair F3+R3d on cDNA obtained from a bulk of 12 days-old plantlets grown in vitro from Col-0, Can-0, Sha, Ang-0 and 9 IP strains carrying either the *Can* (green) or the *Col* (red) allele at the position 28,958,437bp. The red arrows indicate the expected size in Col-0.

**Supplemental figure 10:**
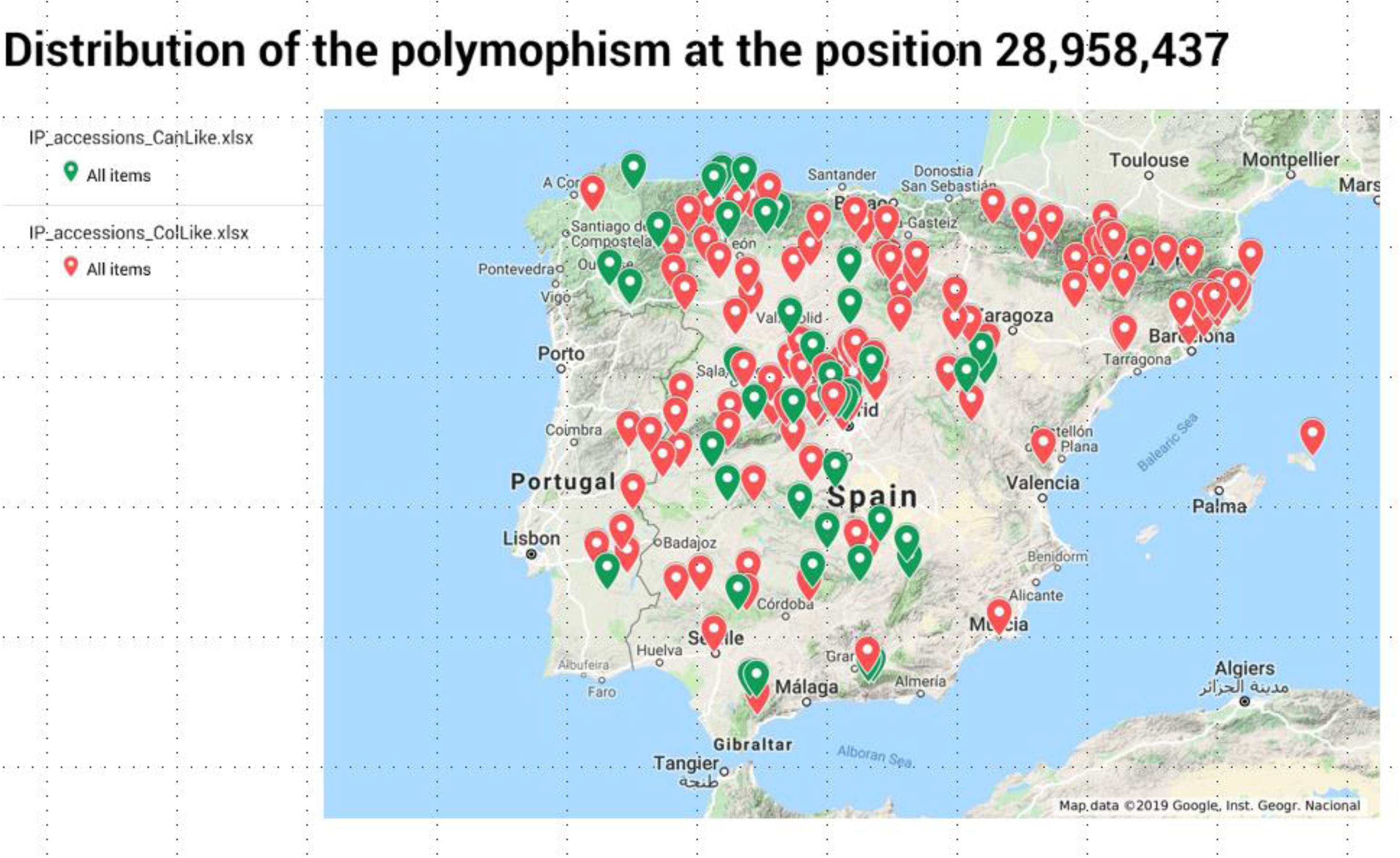
Geographical origin of the IP strains. Map obtained using googlemap showing the location of the IP strains used in this study for the correlation analysis. Location points are respectively green (red) according to the *Can* (*Col*) allele at the position 28,958,437bp on chromosome 1.

## REFERENCES

Ågren, J. and Schemske, D.W. (2012). Reciprocal transplants demonstrate strong adaptive differentiation of the model organism Arabidopsis thaliana in its native range. New Phytol. 194: 1112–1122.

Allen Orr, H. (2006). Adaptation and the Cost of Complexity. Evolution (N. Y). 54: 13.

Alonso-Blanco, C. et al. (2016). 1,135 Genomes Reveal the Global Pattern of Polymorphism in Arabidopsis thaliana. Cell 166: 481–491.

Angert, A.L., Huxman, T.E., Barron-Gafford, G.A., Gerst, K.L., and Venable, D.L. (2007). Linking growth strategies to long-term population dynamics in a guild of desert annuals. J. Ecol. 95: 321–331.

Arends, D., Prins, P., Jansen, R.C., and Broman, K.W. (2010). R/qtl: High-throughput multiple QTL mapping. Bioinformatics 26: 2990–2992.

Bac-Molenaar, J.A., Granier, C., Keurentjes, J.J.B., and Vreugdenhil, D. (2016). Genome-wide association mapping of time-dependent growth responses to moderate drought stress in Arabidopsis. Plant Cell Environ. 39: 88–102.

Balasubramanian, S., Sureshkumar, S., Lempe, J., and Weigel, D. (2006). Potent induction of Arabidopsis thaliana flowering by elevated growth temperature. PLoS Genet. 2: 0980–0989.

Baxter, I., Brazelton, J.N., Yu, D., Huang, Y.S., Lahner, B., Yakubova, E., Li, Y., Bergelson, J., Borevitz, J.O., Nordborg, M., Vitek, O., and Salt, D.E. (2010). A coastal cline in sodium accumulation in Arabidopsis thaliana is driven by natural variation of the sodium transporter AtHKT1;1. PLoS Genet. 6.

Blümel, M., Dally, N., and Jung, C. (2015). Flowering time regulation in crops-what did we learn from Arabidopsis? Curr. Opin. Biotechnol. 32: 121–129.

Brachi, B., Meyer, C.G., Villoutreix, R., Platt, A., Morton, T.C., Roux, F., and Bergelson, J. (2015). Coselected genes determine adaptive variation in herbivore resistance throughout the native range of *Arabidopsis thaliana*. Proc. Natl. Acad. Sci. 112: 4032–4037.

Bradshaw Jr, H. and Schemske, D. (2003). Allele substitution at a flower colour locus produces a pollinator shift in monkeyflower. Nature 426.

Brennan, A.C., Méndez-Vigo, B., Haddioui, A., Martínez-Zapater, J.M., Picó, F.X., and Alonso-Blanco, C. (2014). The genetic structure of Arabidopsis thaliana in the south-western Mediterranean range reveals a shared history between North Africa and southern Europe. BMC Plant Biol. 14.

Broman, K.W., Wu, H., Sen, S., and Churchill, G.A. (2003). R/qtl: QTL mapping in experimental crosses. Bioinformatics 19: 889–890.

Campitelli, B.E., Des Marais, D.L., and Juenger, T.E. (2016). Ecological interactions and the fitness effect of water-use efficiency: Competition and drought alter the impact of natural MPK12 alleles in Arabidopsis. Ecol. Lett. 19: 424–434.

Capovilla, G., Symeonidi, E., Wu, R., and Schmid, M. (2017). Contribution of major FLM isoforms to temperature-dependent flowering in Arabidopsis thaliana. J. Exp. Bot. 68: 5117–5127.

Chen, M. and Penfield, S. (2018). and Flowering Time. Science (80-.). 12: 1014–1017.

Chen, Q., Han, Y., Liu, H., Wang, X., Sun, J., Zhao, B., Li, W., Tian, J., Liang, Y., Yan, J., Yang, X., and Tian, F. (2018). Genome-Wide Association Analyses Reveal the Importance of Alternative Splicing in Diversifying Gene Function and Regulating Phenotypic Variation in Maize. Plant Cell 30: pc.00109.2018.

Chiang, G.C.K., Barua, D., Kramer, E.M., Amasino, R.M., and Donohue, K. (2009). Major flowering time gene, *FLOWERING LOCUS C*, regulates seed germination in *Arabidopsis thaliana*. Proc. Natl. Acad. Sci. 106: 11661–11666.

Cho, L.H., Yoon, J., and An, G. (2017). The control of flowering time by environmental factors. Plant J. 90: 708–719.

Darwin C. (1859) On the Origin of Species by Means of Natural Selection, or the Preservation of Favoured Races in the Struggle for Life. John Murray, London, UK.

Deng, W., Ying, H., Helliwell, C.A., Taylor, J.M., Peacock, W.J., and Dennis, E.S. (2011). FLOWERING LOCUS C (FLC) regulates development pathways throughout the life cycle of Arabidopsis. Proc. Natl. Acad. Sci. 108: 6680–6685.

Díaz, S. et al. (2016). The global spectrum of plant form and function. Nature 529: 167–171.

Donovan, L.A., Maherali, H., Caruso, C.M., Huber, H., and de Kroon, H. (2011). The evolution of the worldwide leaf economics spectrum. Trends Ecol. Evol. 26: 88–95.

Fisher, R. A. 1930. The genetical theory of natural selection. Oxford Univ. Press, Oxford, U.K.

Fisher, R. A. The Genetical Theory of Natural Selection (Dover, New York, 1958).

Frachon, L. et al. (2017). Intermediate degrees of synergistic pleiotropy drive adaptive evolution in ecological time. Nat. Ecol. Evol. 1: 1551–1561.

Frachon, L., Bartoli, C., Carrère, S., Bouchez, O., Chaubet, A., Gautier, M., Roby, D., and Roux, F. (2018). A Genomic Map of Climate Adaptation in Arabidopsis thaliana at a Micro-Geographic Scale. Front. Plant Sci. 9: 1–15.

Fulgione, A. and Hancock, A.M. (2018). Archaic lineages broaden our view on the history of Arabidopsis thaliana. New Phytol. 219: 1194–1198.

Fulgione, A., Koornneef, M., Roux, F., Hermisson, J., and Hancock, A.M. (2018). Madeiran arabidopsis thaliana reveals ancient long-range colonization and clarifies demography in eurasia. Mol. Biol. Evol. 35: 564–574.

Fusari, C.M., Kooke, R., Lauxmann, M.A., Annunziata, M.G., Enke, B., Hoehne, M., Krohn, N., Becker, F.F.M., Schlereth, A., Sulpice, R., Stitt, M., and Keurentjes, J.J.B. (2017). Genome-Wide Association Mapping Reveals That Specific and Pleiotropic Regulatory Mechanisms Fine-Tune Central Metabolism and Growth in Arabidopsis. Plant Cell 29: 2349–2373.

Garnier, E., Navas, M.-L., and Grigulis, K. (2017). Plant Functional Diversity. Organism Traits, Community Structure and Ecosystem Properties. Austral Ecol. 42: e21–e22.

Granier, C., Aguirrezabal, L., Chenu, K., Cookson, S.J., Dauzat, M., Hamard, P., Thioux, J.-J., Bouchier-Combaud, S., Lebaudy, A., Muller, B., Simonneau, T., and Tardieu, F. (2006). PHENOPSIS, an automated platform for reproducible phenotyping of plant responses to soil water deficit in. New Phytol. 169: 623–635.

Grime, J.P. (1977). Evidence for the Existence of Three Primary Strategies in Plants and Its Relevance to Ecological and Evolutionary Theory. Am. Nat. 111: 1169–1194.

Gujas, B., Alonso-Blanco, C., and Hardtke, C.S. (2012). Natural arabidopsis brx loss-of-function alleles confer root adaptation to acidic soil. Curr. Biol. 22: 1962–1968.

Hsu, C.W., Lo, C.Y., and Lee, C.R. (2019). On the postglacial spread of human commensal Arabidopsis thaliana: journey to the East. New Phytol. 222: 1447–1457.

Kalyna, M. et al. (2012). Alternative splicing and nonsense-mediated decay modulate expression of important regulatory genes in Arabidopsis. Nucleic Acids Res. 40: 2454–2469.

Kattge, J. et al. (2011). TRY - a global database of plant traits. Glob. Chang. Biol. 17: 2905–2935.

Kesari, R., Lasky, J.R., Villamor, J.G., Des Marais, D.L., Chen, Y.-J.C., Liu, T.-W., Lin, W., Juenger, T.E., and Verslues, P.E. (2012). Intron-mediated alternative splicing of Arabidopsis P5CS1 and its association with natural variation in proline and climate adaptation. Proc. Natl. Acad. Sci. 109: 9197–9202.

Kooyers, N.J. (2015). The evolution of drought escape and avoidance in natural herbaceous populations. Plant Sci. 234: 155–162.

Kronholm, I., Pico, X., Goudet, J., Alonso-blanco, C., and Meaux, J. De (2012). ARABIDOPSIS THALIANA: LOCAL ADAPTATION AT THE SEED DORMANCY QTL DOG1. Evolution (N. Y).: 2287–2302.

Lee, C.R., Svardal, H., Farlow, A., Exposito-Alonso, M., Ding, W., Novikova, P., Alonso-Blanco, C., Weigel, D., and Nordborg, M. (2017). On the post-glacial spread of human commensal Arabidopsis thaliana. Nat. Commun. 8: 1–12.

Lee, H., Suh, S.S., Park, E., Cho, E., Ahn, J.H., Kim, S.G., Lee, J.S., Kwon, Y.M., and Lee, I. (2000). The AGAMOUS-lIKE 20 MADS domain protein integrates floral inductive pathways in Arabidopsis. Genes Dev. 14: 2366–2376.

Li, P., Tao, Z., and Dean, C. (2015). Phenotypic evolution through variation in splicing of the noncoding RNA COOLAIR. Genes Dev. 29: 696–701.

Liu, L., Adrian, J., Pankin, A., Hu, J., Dong, X., Von Korff, M., and Turck, F. (2014). Induced and natural variation of promoter length modulates the photoperiodic response of FLOWERING LOCUS T. Nat. Commun. 5.

Loudet, O., Gaudon, V., Trubuil, A., and Daniel-Vedele, F. (2005). Quantitative trait loci controlling root growth and architecture in Arabidopsis thaliana confirmed by heterogeneous inbred family. Theor. Appl. Genet. 110: 742–753.

Loudet, O., Saliba-Colombani, V., Camilleri, C., Calenge, F., Gaudon, V., Koprivova, A., North, K.A., Kopriva, S., and Daniel-Vedele, F. (2007). Natural variation for sulfate content in Arabidopsis thaliana is highly controlled by APR2. Nat. Genet. 39: 896–900.

Lovell, J.T., Juenger, T.E., Michaels, S.D., Lasky, J.R., Platt, A., Richards, J.H., Yu, X., Easlon, H.M., Sen, S., and McKay, J.K. (2013). Pleiotropy of FRIGIDA enhances the potential for multivariate adaptation. Proc. R. Soc. B Biol. Sci. 280: 20131043–20131043.

Low, K., Lim, C., Ko, H., and Edery, I. (2008). Natural variation in the splice site strength of a clock gene and species-specific thermal adaptation. Neuron 377: 364–377.

Ludlow, M.M. (1989). Strategies of response to water stress. In: Eds K.H. Kreeb; H. Richter; T.M. Hinckley, editor/s. Structural and Functional Responses to Environmental Stresses, SPB Academic, The Hague (1989), pp. 269–281.

Lutz, U., Nussbaumer, T., Spannagl, M., Diener, J., Mayer, K.F.X., and Schwechheimer, C. (2017). Natural haplotypes of FLM non-coding sequences fine-tune flowering time in ambient spring temperatures in arabidopsis. Elife 6: 1–22.

Lutz, U., Posé, D., Pfeifer, M., Gundlach, H., Hagmann, J., Wang, C., Weigel, D., Mayer, K.F.X., Schmid, M., and Schwechheimer, C. (2015). Modulation of Ambient Temperature-Dependent Flowering in Arabidopsis thaliana by Natural Variation of FLOWERING LOCUS M. PLoS Genet. 11: 1–26.

Mackay, T.F.C. (2004). Complementing complexity. Nat. Genet. 36: 1145–1147.

Mandadi, K.K. and Scholthof, K.-B.G. (2015). Genome-Wide Analysis of Alternative Splicing Landscapes Modulated during Plant-Virus Interactions in Brachypodium distachyon. Plant Cell Online 27: 71–85.

Marchadier, E., Hanemian, M., Tisné, S., Bach, L., Bazakos, C., Gilbault, E., Haddadi, P., Virlouvet, L., and Loudet, O. (2019). The complex genetic architecture of shoot growth natural variation in Arabidopsis thaliana. PLoS Genet. 15: e1007954.

Masle, J., Gilmore, S.R., and Farquhar, G.D. (2005). The ERECTA gene regulates plant transpiration efficiency in Arabidopsis. Nature 436: 866–870.

McKay, J.K., Richards, J.H., and Mitchell-Olds, T. (2003). Genetics of drought adaptation in Arabidopsis thaliana: I. Pleiotropy contributes to genetic correlations among ecological traits. Mol. Ecol. 12: 1137–1151.

Mendez-Vigo, B., Pico, F.X., Ramiro, M., Martinez-Zapater, J.M., and Alonso-Blanco, C. (2011). Altitudinal and Climatic Adaptation Is Mediated by Flowering Traits and FRI, FLC, and PHYC Genes in Arabidopsis. Plant Physiol. 157: 1942–1955.

Michaels, S.D. and Amasino, R.M. (1999). FLOWERING LOCUS C encodes a novel MADS domain protein that acts as a repressor of flowering. Plant Cell 11: 949–56.

Monroe, J.G., Powell, T., Price, N., Mullen, J.L., Howard, A., Evans, K., Lovell, J.T., and McKay, J.K. (2018). Drought adaptation in Arabidopsis thaliana by extensive genetic loss-of-function. Elife 7: 1–18.

Orr, H.A. (2005). The genetic theory of adaptation: A brief history. Nat. Rev. Genet. 6: 119–127.

Paaby, A.B. and Rockman, M. V. (2013). The many faces of pleiotropy. Trends Genet. 29: 66–73.

Park, E., Pan, Z., Zhang, Z., Lin, L., and Xing, Y. (2018). The Expanding Landscape of Alternative Splicing Variation in Human Populations. Am. J. Hum. Genet. 102: 11–26.

Van De Peer, Y., Mizrachi, E., and Marchal, K. (2017). The evolutionary significance of polyploidy. Nat. Rev. Genet. 18: 411–424.

Poormohammad Kiani, S., Trontin, C., Andreatta, M., Simon, M., Robert, T., Salt, D.E., and Loudet, O. (2012). Allelic heterogeneity and trade-off shape natural variation for response to soil micronutrient. PLoS Genet. 8: 3–8.

Posé, D., Verhage, L., Ott, F., Yant, L., Mathieu, J., Angenent, G.C., Immink, R.G.H., and Schmid, M. (2013). Temperature-dependent regulation of flowering by antagonistic FLM variants. Nature 503: 414–417.

Postma, F.M. and Ågren, J. (2016). Early life stages contribute strongly to local adaptation in Arabidopsis thaliana. Proc. Natl. Acad. Sci. 113: 7590–7595.

Prasad, K.V.S.K. et al. (2012). A gain-of-function polymorphism controlling complex traits and fitness in nature. Science (80-.). 337: 1081–1084.

Pritchard, J.K., Pickrell, J.K., and Coop, G. (2010). The Genetics of Human Adaptation: Hard Sweeps, Soft Sweeps, and Polygenic Adaptation. Curr. Biol. 20: R208–R215.

Reddy, A.S.N., Marquez, Y., Kalyna, M., and Barta, A. (2013). Complexity of the Alternative Splicing Landscape in Plants. Plant Cell 25: 3657–3683.

Reich, P.B. (2014). The world-wide “fast-slow” plant economics spectrum: A traits manifesto. J. Ecol. 102: 275–301.

Sartori, K.F.R., Vasseur, F., Violle, C., Baron, E., Gerard, M., Rowe, N., Ayala-Garay, O., Christophe, A., De Jalon, L.G., and Masclef, D. (2018). Leaf economics guides slow-fast adaptation across the geographic range of *A. thaliana*. bioRxiv: 487066.

Sass, L., Majer, P., and Hideg, É. (2012). Leaf Hue Measurements: A High-Throughput Screening of Chlorophyll Content. In High-Throughput Phenotyping in Plants: Methods and Protocols, J. Normanly, ed (Humana Press: Totowa, NJ), pp. 61–69.

Scortecci, K.C., Michaels, S.D., and Amasino, R.M. (2001). Identification of a MADS-box gene, FLOWERING LOCUS M, that represses flowering. Plant J. 26: 229–236.

Sharbel, T.F., Haubold, B., and Mitchell-Olds, T. (2000). Genetic isolation by distance in Arabidopsis thaliana: biogeography and postglacial colonization of Europe. Mol. Ecol. 9: 2109–2118.

Sheehan, H., Moser, M., Klahre, U., Esfeld, K., Dell’olivo, A., Mandel, T., Metzger, S., Vandenbussche, M., Freitas, L., and Kuhlemeier, C. (2016). MYB-FL controls gain and loss of floral UV absorbance, a key trait affecting pollinator preference and reproductive isolation. Nat. Genet. 48: 159–166.

Shen, Y., Zhou, Z., Wang, Z., Li, W., Fang, C., Wu, M., Ma, Y., Liu, T., Kong, L.-A., Peng, D.-L., and Tian, Z. (2014). Global Dissection of Alternative Splicing in Paleopolyploid Soybean. Plant Cell 26: 996–1008.

Simon, M., Loudet, O., Durand, S., Bérard, A., Brunel, D., Sennesal, F.X., Durand-Tardif, M., Pelletier, G., and Camilleri, C. (2008). Quantitative trait loci mapping in five new large recombinant inbred line populations of Arabidopsis thaliana genotyped with consensus single-nucleotide polymorphism markers. Genetics 178: 2253–2264.

Stinchcombe, J.R., Weinig, C., Ungerer, M., Olsen, K.M., Mays, C., Halldorsdottir, S.S., Purugganan, M.D., and Schmitt, J. (2004). A latitudinal cline in flowering time in Arabidopsis thaliana modulated by the flowering time gene FRIGIDA. Proc. Natl. Acad. Sci. 101: 4712–4717.

Sureshkumar, S., Dent, C., Seleznev, A., Tasset, C., and Balasubramanian, S. (2016). Nonsense-mediated mRNA decay modulates FLM-dependent thermosensory flowering response in Arabidopsis. Nat. Plants 2: 1–16.

Thatcher, S.R., Zhou, W., Leonard, A., Wang, B.-B., Beatty, M., Zastrow-Hayes, G., Zhao, X., Baumgarten, A., and Li, B. (2014). Genome-Wide Analysis of Alternative Splicing in Zea mays: Landscape and Genetic Regulation. Plant Cell 26: 3472–3487.

Tisné, S. et al. (2013). Phenoscope: An automated large-scale phenotyping platform offering high spatial homogeneity. Plant J. 74: 534–544.

Tisné, S., Schmalenbach, I., Reymond, M., Dauzat, M., Pervent, M., Vile, D., and Granier, C. (2010). Keep on growing under drought: Genetic and developmental bases of the response of rosette area using a recombinant inbred line population. Plant, Cell Environ. 33: 1875–1887.

Trontin, C., Kiani, S., Corwin, J.A., Hématy, K., Yansouni, J., Kliebenstein, D.J., and Loudet, O. (2014). A pair of receptor-like kinases is responsible for natural variation in shoot growth response to mannitol treatment in Arabidopsis thaliana. Plant J. 78: 121–133.

Vasseur, F., Bontpart, T., Dauzat, M., Granier, C., and Vile, D. (2014). Multivariate genetic analysis of plant responses to water deficit and high temperature revealed contrasting adaptive strategies. J. Exp. Bot. 65: 6457–6469.

Vasseur, F., Sartori, K., Baron, E., Fort, F., Kazakou, E., Segrestin, J., Garnier, E., Vile, D., and Violle, C. (2018). Climate as a driver of adaptive variations in ecological strategies in Arabidopsis thaliana. Ann. Bot.: 1–11.

Vasseur, F., Violle, C., Enquist, B.J., Granier, C., and Vile, D. (2012). A common genetic basis to the origin of the leaf economics spectrum and metabolic scaling allometry. Ecol. Lett. 15: 1149–1157.

Vidigal, D.S., Marques, A.C.S.S., Willems, L.A.J., Buijs, G., Méndez-Vigo, B., Hilhorst, H.W.M., Bentsink, L., Picó, F.X., and Alonso-Blanco, C. (2016). Altitudinal and climatic associations of seed dormancy and flowering traits evidence adaptation of annual life cycle timing in Arabidopsis thaliana. Plant Cell Environ. 39: 1737–1748.

Violle, C., Navas, M.L., Vile, D., Kazakou, E., Fortunel, C., Hummel, I., and Garnier, E. (2007). Let the concept of trait be functional! Oikos 116: 882–892.

Vlad, D., Rappaport, F., Simon, M., and Loudet, O. (2010). Gene transposition causing natural variation for growth in Arabidopsis thaliana. PLoS Genet. 6: 21.

Wagner, G.P., Kenney-Hunt, J.P., Pavlicev, M., Peck, J.R., Waxman, D., and Cheverud, J.M. (2008). Pleiotropic scaling of gene effects and the “cost of complexity.” Nature 452: 470–472.

Wagner, G.P., Pavlicev, M., and Cheverud, J.M. (2007). The road to modularity. Nat. Rev. Genet. 8: 921–931.

Wagner, G.P. and Zhang, J. (2011). The pleiotropic structure of the genotype-phenotype map: The evolvability of complex organisms. Nat. Rev. Genet. 12: 204–213.

Wang, Z., Liao, B.-Y., and Zhang, J. (2010). Genomic patterns of pleiotropy and the evolution of complexity. Proc. Natl. Acad. Sci. 107: 18034–18039.

Whittaker, C. and Dean, C. (2017). The FLC Locus: A Platform for Discoveries in Epigenetics and Adaptation. Annu. Rev. Cell Dev. Biol. 33: 555–575.

Willmann, M.R. and Poethig, R.S. (2011). The effect of the floral repressor FLC on the timing and progression of vegetative phase change in Arabidopsis. Development 138: 677–685.

Wright, I.J. et al. (2004). The worldwide leaf economics spectrum. Nature 428: 821–827.

Wright, I.J. (2004). The worldwide leaf economics spectrum. Nature 428: 821–827.

Wu, W. et al. (2017). A single-nucleotide polymorphism causes smaller grain size and loss of seed shattering during African rice domestication. Nat. Plants 3.

Xu, Y.-C., Niu, X.-M., Li, X.-X., He, W., Chen, J.-F., Zou, Y.-P., Wu, Q., Zhang, Y.E., Busch, W., and Guo, Y.-L. (2019). Adaptation and phenotypic diversification through loss-of-function mutations in Arabidopsis protein-coding genes. Plant Cell 31: pc.00791.2018.

Zhang, G. et al. (2010). Deep RNA sequencing at single base-pair resolution reveals high complexity of the rice transcriptome. Genome Res. 20: 646–54.

